# Conservation networks do not match the ecological requirements of amphibians

**DOI:** 10.1101/2022.07.18.500425

**Authors:** Matutini Florence, Jacques Baudry, Marie-Josée Fortin, Guillaume Pain, Joséphine Pithon

## Abstract

Amphibians are among the most threatened taxa as they are highly sensitive to habitat degradation and fragmentation. They are considered as model species to evaluate habitats quality in agricultural landscapes. In France, all amphibian species have a protected status requiring recovery plans for their conservation. Conservation networks combining protected areas and green infrastructure can help the maintenance of their habitats while favouring their movement in fragmented landscapes such as farmlands. Yet, assessing the effectiveness of conservation networks is challenging.
Here, we compared the ecological requirements of amphibian species with existing conservation network coverage in a human-dominated region of western France. First, we mapped suitable habitat distributions for nine species of amphibian with varying ecological requirements and mobility. Second, we used stacking species distribution modelling (SSDM) to produce multi-species habitat suitability maps. Then, to identify spatial continuity in suitable habitats at the regional scale, we defined species and multi-species core habitats to perform a connectivity analysis using Circuitscape theory. Finally, we compared different suitability maps with existing conservation networks to assess conservation coverage and efficiency.
We highlighted a mismatch between the most suitable amphibian habitats at the regional scale and the conservation network, both for common species and for species of high conservation concern. We also found two bottlenecks between areas of suitable habitat which might be crucial for population movements induced by global change, especially for species associated with hedgerow mosaic landscapes. These bottlenecks were not covered by any form of protection and are located in an intensive farmland context.
*Synthesis and applications* - We advocate the need to better integrate agricultural landscape mosaics into species conservation planning as well as to protect and promote agroecological practices suitable for biodiversity, including mixed and extensive livestock farming. We also emphasize the importance of interacting landscape elements of green infrastructure for amphibian conservation and the need for these to be effectively considered in land-use planning policies.

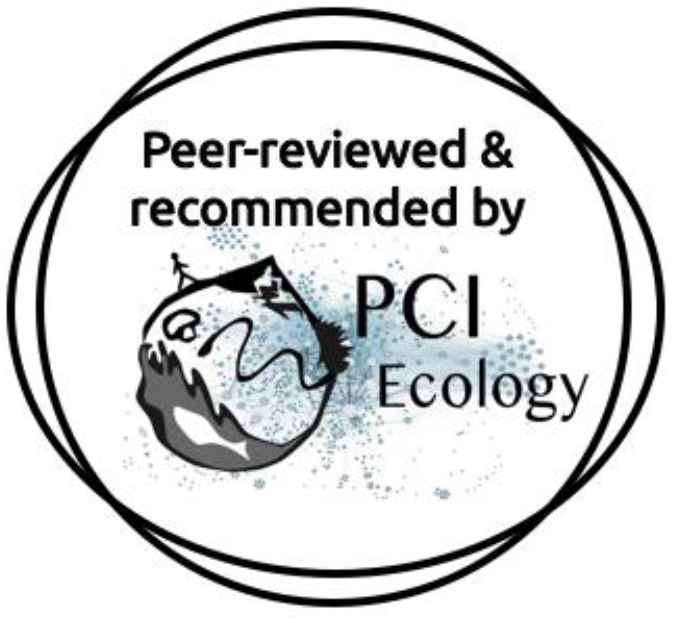

PCI recommendation : https://ecology.peercommunityin.org/articles/rec?id=504

## 1 Introduction

The biodiversity decline is still accelerating across the world affecting both scarce and common species (Dirzo et al., 2014; Gaston & Fuller, 2008) especially in agroecosystems (Kleijn et al., 2011; Seibold et al., 2019). To palliate this decline, the establishment of protected areas (PAs) is a relevant conservation tool in a human-dominated world (Rodrigues & Cazalis, 2020). However, PAs alone are not sufficient to protect biodiversity and they should be part of an ensemble of management and planning strategies aiming to create functional conservation networks. Green infrastructure (GI) (also called Green and Blue infrastructure in France) has been proposed as a way to address this issue (Chatzimentor et al., 2020; Salomaa et al., 2017). Such initiatives aim to maintain a coherent and functional network of suitable habitats and to improve biodiversity integration into land-use planning. GI design is based on the ecological network concept (Opdam et al., 2006) emphasizing the complementarity of core habitat patches and ecological corridors at different spatial and temporal scales (Opdam et al., 2006; Salomaa et al., 2017).

While conservation networks are crucial for biodiversity conservation, they might have some weaknesses in their design that need to be addressed. For example, ecological network design from landscape connectivity modelling poorly integrates biological data which may be problematic in operational contexts (Foltête et al., 2020). In France, GI has mostly been defined based on spatial analysis of land-cover data without considering species data. It is based on a simplistic classification of ecosystems reduced to broad categories of land cover data (i.e. different layers for forest or wetlands separately) with a poor consideration of habitat complementation. In contrast, PAs have usually been defined according to the needs of focal endangered species and/or habitats, often charismatic species such as birds and endangered plants, which might be poorly representative of species-habitat relations and ecosystem processes (Rodrigues et al., 2004).

Evaluating the efficiency of conservation networks (here PAs and GI) is a real challenge (e.g., Rodrigues & Cazalis, 2020). Gap analysis and conservation prioritization are common approaches for a first-step assessment of the effectiveness of a conservation network and to inform decisions for conservation measures (Jennings 2000; Moilanen et al. 2005). To conduct a gap analysis, outputs of spatial modelling are particularly useful (Ahmadi et al., 2020). Species distribution models (SDMs) are commonly used to make spatially explicit predictions of the suitable environment of a given species from biological data (Guisan & Zimmermann, 2000) and multi-specific approaches have recently been developed (Scherrer et al., 2018; Thuiller et al., 2015). In addition, recent methods integrate both graph theory and SDMs for ecological network modelling (Duflot et al., 2018; Godet & Clauzel, 2020). These multi-species and integrative approaches could be relevant for gap analyses and spatial prioritization but remain underused (Foltête et al., 2020).

In western Europe, traditional hedgerow landscapes are of high conservation concern for numerous taxa such as plants, amphibians, birds, bats, and arthropods (Baudry et al., 2000; Boissinot et al., 2019). In France, these rural landscapes are associated with dense hedgerow networks interconnected with a mosaic of small, interlocked patches of pastures, cultivated fields and bushes with high pond density (Burel & Baudry, 1995). They have been greatly impacted by agricultural intensification and have declined from 40 % to 80 % in Europe (Bazin & Schmutz, 1994). Amphibians are good candidates to assess the ecological quality of human-dominated landscape mosaics and to question the efficiency of existing conservation networks. Indeed, they are among the most threatened taxa with rapid and widespread population declines in both natural and modified habitats (Stuart et al., 2004). They have a bi-phasic lifecycle with different ecological requirements inducing seasonal migrations between habitat patches (Sinsch, 2014). Their poor mobility and their permeable skin increase their sensitivity to anthropogenic disturbances, pollution and ecological fragmentation induced by urbanization and intensive agriculture (Cushman, 2006; Hamer & McDonnell, 2008; Stuart et al., 2004). Hence, some authors consider amphibians as good ecological indicators of general environmental health (Collins and Storfer 2003; Díaz-García et al. 2017 but see Sewell & Griffiths, 2009). In addition, amphibians have been the target of several citizen science programs in different countries around the world (De Solla et al., 2005; Petrovan & Schmidt, 2016) leading to a large amount of available data over wide areas which could be relevant for conservation tools assessment (Snäll et al., 2016).

The aim of this study is to propose a method to assess the ecological quality of agricultural mosaics landscapes from species data and to conduct a gap analysis on existing conservation networks. In a human-dominated region of western France with traditional hedgerow landscapes of high conservation concern, we compared the predicted habitat requirements of amphibian species with existing conservation network coverage (of both protected areas and green infrastructure). Specifically, we: (1) used single habitat suitability maps from Matutini et al., (2021) and produced multi-species habitat suitability maps for amphibians with differing ecological requirements and mobility (2) compared these habitat suitability maps with existing conservation networks (PAs and GI), and (3) identified gaps in conservation coverage which could be a priority for new potential conservation areas. Additional analyses on all the species in the region were carried out to discuss the potential generalization of the results obtained on rare species.

## 2 Material and Methods

### 2.1 Regional context and studied species

#### Ecological context

The study area is located in a region of western France (Pays-de-la-Loire) covering 32,082 km^2^. Intensification of agricultural practices has greatly transformed the landscape over the last century from a complex matrix of small fields, meadows and woody elements (i.e. traditional hedgerow landscapes) to more homogeneous landscapes with larger fields. With 20 native species of amphibian (out of 36 native species recorded in metropolitan France), of which 12 are priority species for conservation, the region has considerable responsibility for the preservation of amphibian species and their habitats, including traditional hedgerow landscapes and wetlands.

#### Studied species

Nine amphibian species in the Pays-de-la-Loire region were studied (see Table 1 for species list and conservation issues). Three form a “FOR” group of forest species or species associated with dense hedgerow mosaic landscape (*Triturus marmoratus, Rana temporaria* and *Salamandra* s*alamandra*) and five form a “GEN” group of more generalist species (*Lissotriton helveticus, Bufo spinosus, Hyla arborea, Rana dalmatina* and *Triturus cristatus). Pelodytes punctatus* is a pioneer species mainly associated with disturbed open environments, emblematic of alluvial valleys. Finally, the “ALL” group contained all nine studied species. Species were selected according to data availability (i.e., at least 500 presence after filtering operations according to the number of predictors used for species distribution modelling, see Matutini et al., (2021a). Supplementary information about data, species selection and other species of the region are presented Appendix 1.

**Table 1.** Description of the studied species. RL FR: France red list (2015); RL Region: regional red list of Pays-de-la-Loire (2021); Priority: regional priority level for species conservation from the regional red list from 0 (low) to 3 (very high) defined by Marchadour et al. (2021); Hab. Dir. N2000: habitats directive classification (Natura 2000); Nb presence data: Number of presence-only data were calculated after a 100 m-resolution rasterization and correspond to the number of pixels with at least one observation of the species. In bold, vulnerable or near threatened species at the regional scale. See appendix 1 for species selection and presentation of other species present in the studied area.

|  | Species | RL FR | RL Region | Hab. Dir.<br>N2000 | Nb presence<br>data (Atlas) |
| --- | --- | --- | --- | --- | --- |
| Species | ***Pelodytes punctatus*** | **LC** | **NT** |  | **1714** |
| used for | ***Triturus marmoratus*** | **NT** | **NT** | **IV** | **906** |
| SSDM | ***Rana temporaria*** | **LC** | **VU** | **V** | **848** |
|  | *Lissotriton helveticus* | LC | LC |  | 4580 |
|  | *Bufo spinosus* | / | LC |  | 6071 |
|  | *Hyla arborea* | NT | LC | IV | 4555 |
|  | *Rana dalmatina* | LC | LC | IV | 6118 |
|  | ***Triturus cristatus*** | **NT** | **NT** | **II & IV** | **1040** |
|  | *Salamandra salamandra* | LC | LC |  | 3514 |

#### Biodiversityconservationpolicies

In France, there are different conservation tools including protected areas, green infrastructures, and inventoried sites with recognised biodiversity value but that are not protected (Categories “PA”,”GI” and “INV” respectively in Table 2). We defined five groups of conservation tool based on IUCN categories and protection levels (Table 2 and Fig. 1). For two types of PAs (SCEN and ENS in PA-Group 2), mapped site boundaries were not available unlike other PAs and GI where we have more precise GIS data. As a result, only a point location according to the site centroid was considered in the analysis (see Appendix 2). If one conservation area was overlapped by another, we classed the area into the group with the strongest protection level and therefore all PAs were excluded from GI. French GI is divided into different land cover or landscape classes (Table 2). Each GI layer includes two types of elements: habitat nodes, considered as “biodiversity reservoirs”, and “corridors” ensuring potential continuity between nodes based on geographical criteria.

**Table 2.**
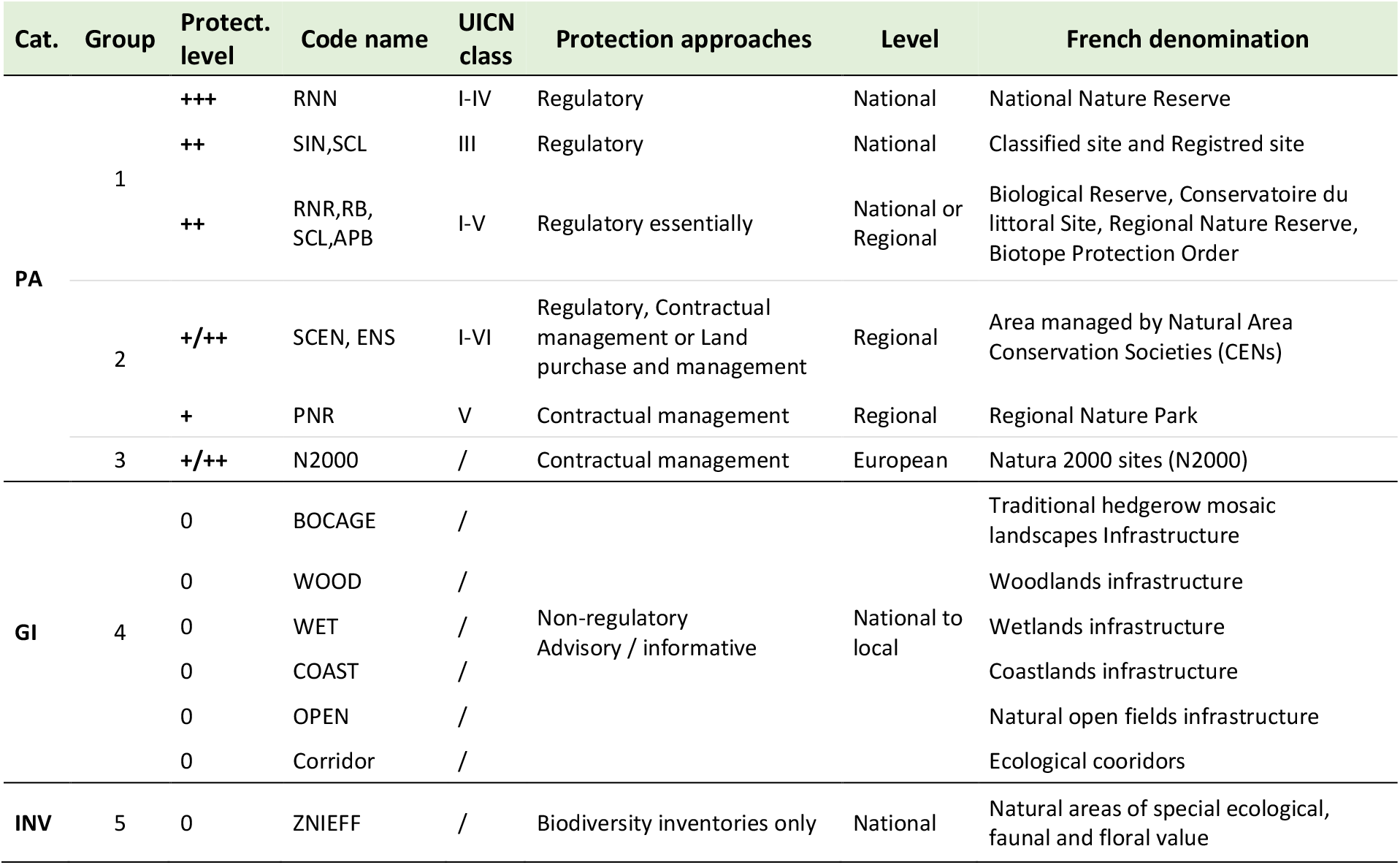
Classification of main conservation areas according to protection levels used in Pays-de-la-Loire (France). PA: protected area; GI: Green infrastructure; INV: area with an inventory of biodiversity and having a strong natural heritage interest but not protected by regulatory tools (ZNIEFF, type I and II). The level of protection varies from 0 (null) to +++ (strong).

**Table 3.** Accuracy of stacked species distribution models for three species groups according to the nine studied amphibian species. Results obtained with external data with 500 permutations. AUC were calculated from predicted multi-species suitability values compared to observed species diversity. Community AUC (AUCc) were obtained with predicted suitability index for each species compared to the presence/absence observations in the assemblage (Scherrer et al. 2020). Individual AUC and SEDI are evaluation metrics obtained by Matutini, Baudry, Fortin, et al. (2021) for each species specific map.

| **Species group** | **R<sup>2</sup>** | **AUC** | **AUCc** | **Min**<br>**individual**<br>**AUC / SEDI** | **Max**<br>**individual**<br>**AUC / SEDI** |
| --- | --- | --- | --- | --- | --- |
| All nine<br>species (*n*=9) | 0.31 | 0.86 | 0.79 | 0.73 / 0.67 | 0.86 / 0.89 |
| Forest species<br>(*n*=3) | 0.23 | 0.76 | 0.96 | 0.78 / 0.68 | 0.86 / 0.84 |
| Generalist<br>species (*n*=5) | 0.30 | 0.87 | 0.86 | 0.73 / 0.67 | 0.81 / 0.89 |

**Fig. 1.**
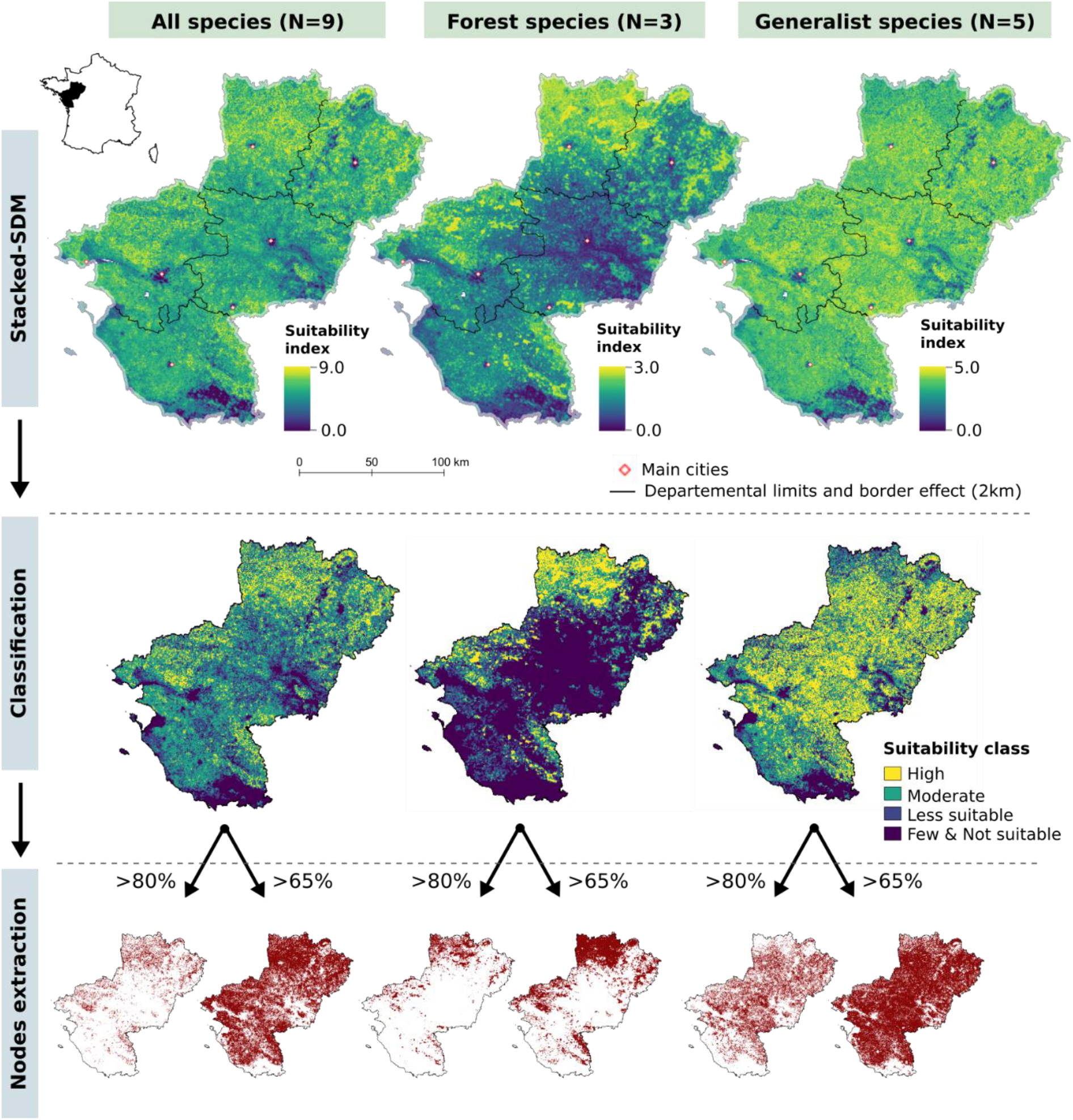
Identification of highly suitable and suitable multi-specific patches of habitat using stacked habitat suitability map for three species groups.

### 2.2 Single and multi-species habitat suitability maps

We used 100 m-resolution habitat suitability maps from Matutini et al., (2021a). This used heterogeneous data from citizen sciences completed by professional surveys to perform and evaluate (with independent dataset) presence-only habitat suitability modelling with functional friction-based and multi-scale predictors (see Matutini et al., 2021 and Matutini et al., 2021b for full details and method tests). Model calibration sets were opportunistic presence-only data from a regional Atlas project with distance-based filtering to reduce spatial autocorrelation coupled with a weighted pseudo-absence selection to reduce sampling bias. SDM were performed combining Random Forest and General additive models. An independent and a standardised detection-nondetection dataset, stratified by model predictions for each species, were used for a robust model evaluation. This dataset included 576 ponds monitored by experts at least 2 nights with a 5-min acoustic survey followed by a visual inspection using halogen light and direct sampling using a fishing net. Data are described Appendix 1.

We produced multi-species suitability maps by compiling individual species maps (Stack-SDM, e.g. Ferrier & Guisan, 2006) using a simple probabilistic stacking method (pSSDM, see Zurell et al., 2019). We summed continuous individual maps without binary “presence-absence” conversion (Calabrese et al., 2014; Scherrer et al., 2020) for three focal species groups: (1) forest species (FOR), (2) generalist species (GEN), and (3) all species (ALL). According to the ecological context of our region (i.e., a hedgerow landscape with small, interlocked landscape elements) for a best integration of mosaic effect for landscape evaluation, we summed species maps at 500 m-resolution after 100 m-pixels aggregation using maximum suitability values (see Appendix 3 for additional information and selection tests). Hence, we compiled single species standard deviation maps to obtain a general representation of levels of confidence in our mapped distributions. To assess the accuracy of the three pSSDMs, we compared multi-species suitability index (i.e., the sum of single species suitability values from 0 to 1) with species richness from independent pond monitoring (Matutini et al., 2021b). We calculated a community-AUC (Scherrer et al., 2020) and *R*^2^ value for each species group with 500 iterations. For each iteration, the evaluation-set was built by randomly selecting the pixels (500m) containing evaluation data respecting a distance of 1km between each pixel (independence) and stratifying the pixels along the suitability gradient of the predicted map (methods tested and validated in Matutini et al., 2021b).

### 2.3 Reclassification and core habitat node definition

For each species and stacked species group, we reclassified the final suitability map into five categories of potential habitat: not suitable (below the ten-percentile value P^10th^ method in Huang et al., 2020), Less suitable (P^10th^-55%), moderately suitable (55%-65%), suitable (65%-80%), and highly suitable (>80%).

### 2.4 Network patterns

To map potential connections between each highly suitable patch (i.e., patch with a suitability index above 80%) at regional scale, we used circuit theory with Circuitscape software (McRae et al., 2016). Circuit theory considered each cell as an electric node connected to neighbouring cells by resistors and defined by a landscape resistance (or conductance) value (McRae et al., 2008). We used highly suitable core habitats as source patches, and a friction map as a measure of conductance (i.e., conductance of each raster 100m cell for movement). This method provides an exploration of potential links between nodes according to landscape permeability combining habitat suitability and connectivity modelling (Ahmadi et al., 2020; Koen et al., 2014).

Habitat suitability could however be a poor proxy of permeability as species may move in unsuitable habitats (Keeley et al., 2017). Hence, the friction map was computed using different friction cost values derived from the multi-species habitat map and landcover data (Keeley et al., 2016, 2017). Therefore, we considered moderately to highly suitable habitat classes (i.e., from 65% to 100%) as permeable features except for pixels crossed by a linear barrier. In less suitable habitats (i.e., under the 65% threshold), we calculated a friction cost based on a 100m-grid analysis performed over 10 m-resolution landscape layers followed by a classification procedure (see Appendix 4). As a precaution, the friction values were defined according to the least mobile species of each group. 100m-resolution were selected to reduce computational power needed for Circuitscape analysis and to consider landscape complementation of fine-scale landscape elements (e.g., ditches, hedgerows and field margins, Pope et al., 2000; Mazerolle, 2005). The cost values of impermeable barriers (i.e., high-density urban area, highspeed train line, highway and dual carriageways) were considered as infinite. Permeable features facilitating movement across barriers (e.g., viaduct, wildlife pass and others, see Table 2) were digitized from aerial photographs. To reduce border effects, we allocated resistance values to a 5km-buffer around the studied region which were randomly selected if land-cover data were not available.

### 2.5 Gap analysis and prioritizing for conservation

We based the gap analysis framework on a method used by Ahmadi et al. (2020) that compared reclassified suitability maps with areas covered by PA. We calculated “total conservation coverage” (i.e., the proportion of suitable habitat overlapping a PA or GI), “conservation efficiency” (i.e., the proportion of PA or GI overlapping suitable habitats) and plotted histograms showing the total distribution of suitability values covered by different types of conservation area. To complete the analysis, we overlaid the existing conservation network with the final regional maps obtained using Circuitscape for each species group to discuss network configuration in relation to potential regional continuities. All spatial analyses were performed using R environment v. 3.5.3 (R Development Core Team, 2019).

## 3 Results

### 3.1 Multi-species maps and core habitat definition

For each group, the AUC varied from 0.76 (forest species) to 0.87 (generalist species) and *R*^2^ from 0.23 (forest species) to 0.31 (nine studied species). The suitable maps are shown in Fig. 1 while associated confidence maps are available Appendix 5. For the nine studied species, highly suitable habitats covered 12% of the region and suitable habitats covered 49% of the region (Fig. 1). Highly suitable habitat for the forest species, covered 11% and suitable habitat covered 17% of the region. Larger amounts of highly suitable and suitable habitats were available to generalist species, covering 33% and 40% of the region respectively.

### 3.2 Protected areas (PAs)

The distribution of protected areas along the habitat suitability gradients modelled for each species or species group (FOR, GEN, ALL) is shown Fig. 2.

**Fig. 2.**
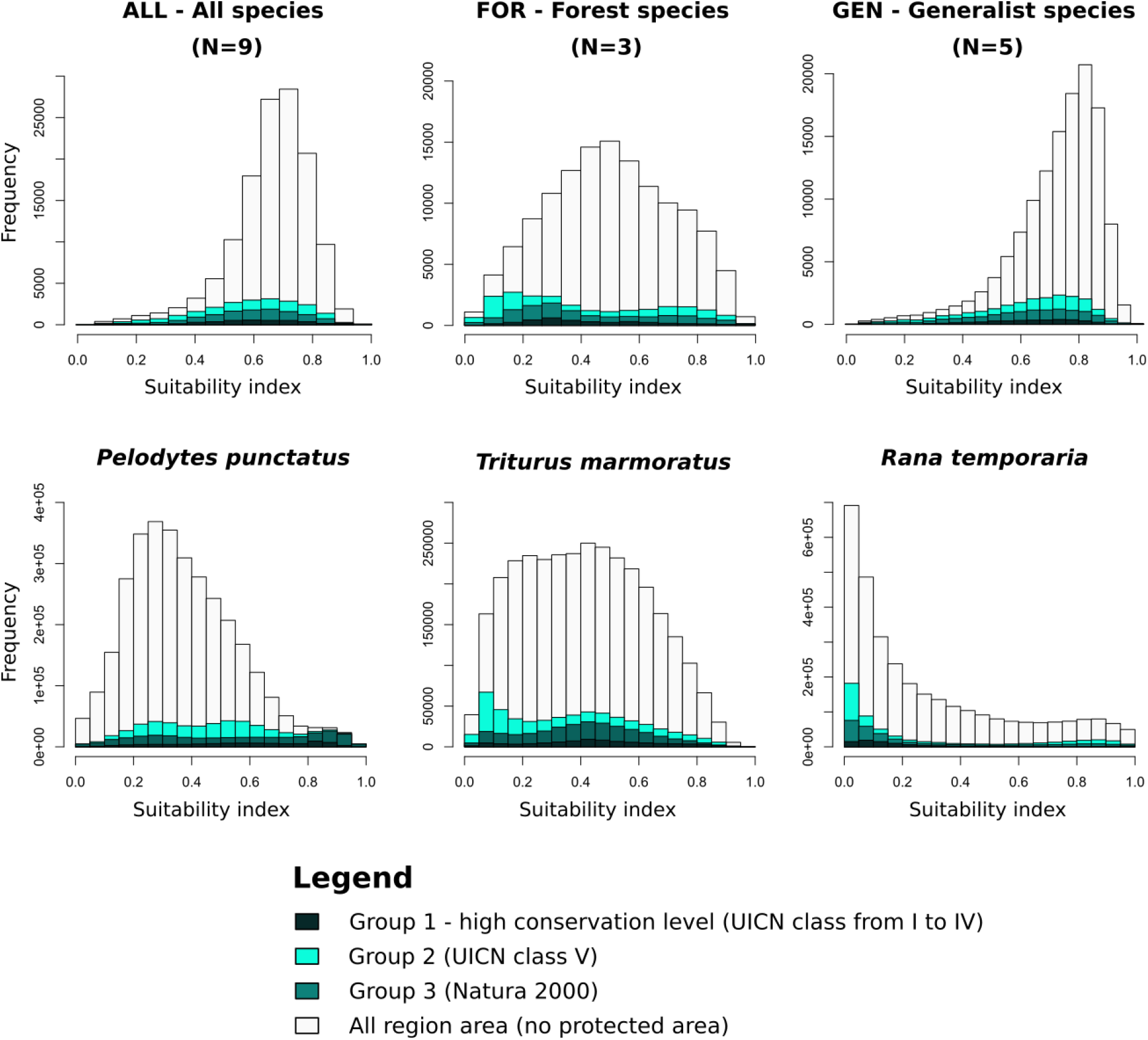
Distribution of pixels along the regional habitat suitability gradient covered or not by protected area. Suitability index (from the habitat suitability map) is calculated for each species and varies from 0 to 1. Suitability index of the three groups (ALL, FOR and GEN) is the sum of the individual species index related to each group (pSSDM) divided by the number of species in order to have a suitability index on the same scale. The frequency corresponds to a number of pixels covered by protected areas from the Group 1, 2, 3 in relation to the whole region area.

#### Conservationcoverage

PAs covered a relatively low proportion of highly suitable habitat: 15%, 19% and 8% for all species, forest species and generalist species respectively (Table 4). The most strictly protected areas protection (PA group 1) covered even lower proportions of suitable habitat: 1% of highly suitable habitat and 2% of suitable habitat for each species group. Similarly, for two of the three species of high conservation concern, PA covered only a relatively low 17% or 26% of highly suitable habitat, for *T. marmoratus* and *R. temporaria* respectively. Only for *P. punctatus* was conservation coverage high: 90% of highly suitable habitat was included in PA. We also found (see Appendix 1) low conservation coverage of PAs for rarer species based on available presence-only data, except for *Pelobates cultripes* with 72% of presence data located in protected areas (of which 37% in PA Group 1).

**Table 4.**
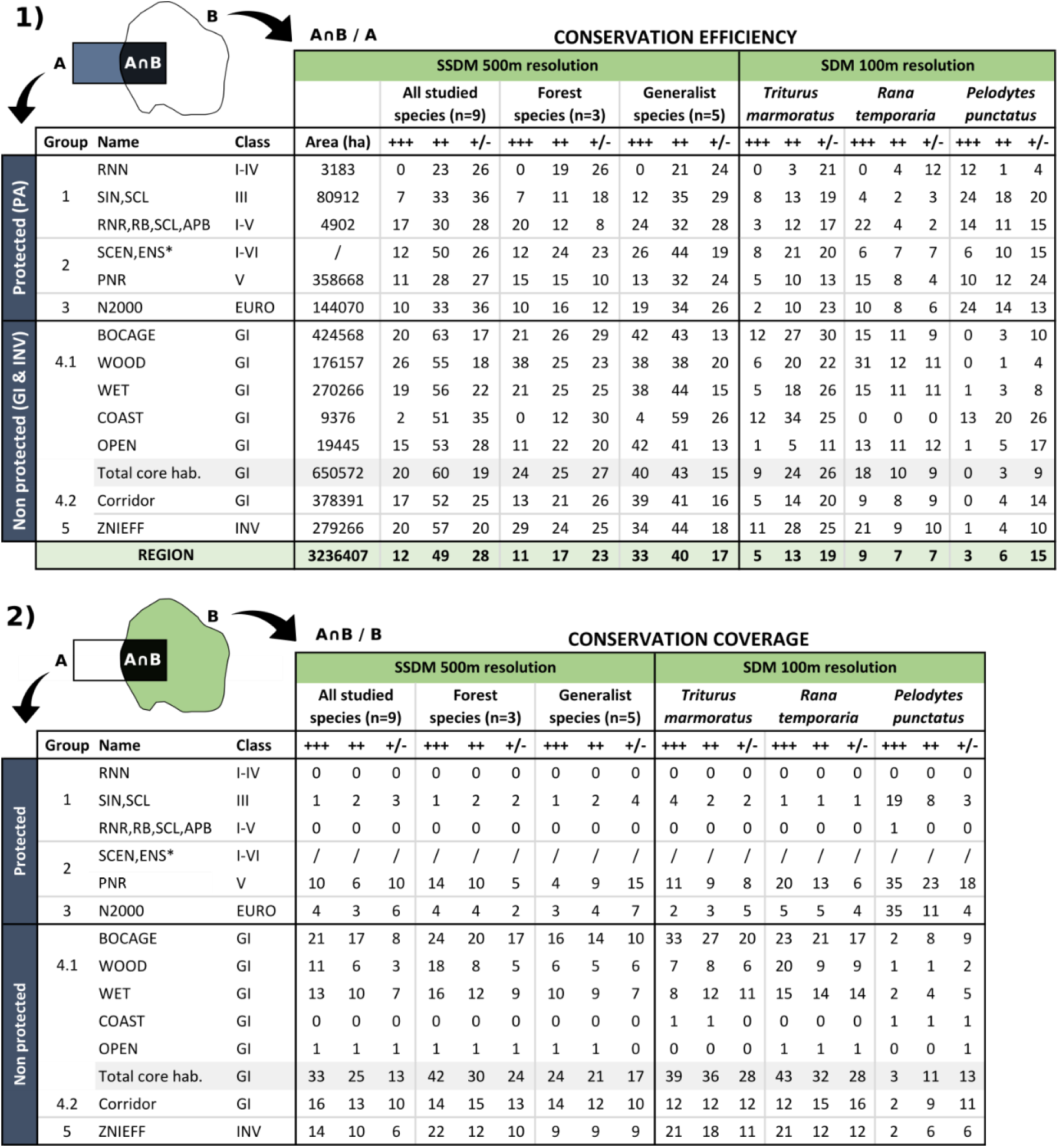
Proportion of the conservation area covered by different habitat suitability classes (1. conservation efficiency) and proportion of different habitat suitability classes covered by the conservation area (2. conservation coverage) for different species and species groups. Classes of conservation area: Protected (groups 1 to 3) – classes I to V derived from the UICN classification; EURO: European conservation network from Natura 2000; Non-protected: GI (Green infrastructure) and INV (inventoried sites define as “high naturalist interest”). Suitability class for each groups or species: “+++” = highly suitable (>0.80) ; “++” = suitable (>0.65) ; +/-= “moderately or not suitable” (<=0.55). See Table 2 for “code names” details. Names with * show sites with point data only (surface area is not available).

#### Conservationefficiency

Among PA-Group 1, no sites with the highest level of protection (names “RNN”, see Table 2) contained highly suitable area for any species group or individual forest species except *P. punctatus* (12% of RNN areas contain high suitable habitat) (Table 4). Other PA-Group 1 protection levels were covered by 7 to 24% of multi-species core area and by 3 to 24% of species-specific highly suitable habitat, with a higher proportion for *P. punctatus*. Regional Park (PA-Group 2) and Natura 2000 network (PA-Group 3) included less than 16% multi-species core area and lower proportions for the two-forest species individually. For SCEN and ENS (in PA-Group 2) 26%, 12% and 8% of these sites were located in highly suitable habitats for generalists, forest species and all nine species respectively. Overall, PAs contain a higher proportion of highly suitable habitat for *P. punctatus* than for other species.

### 3.3 Inventoried sites (INV)

Inventoried sites with known ecological value but no legal protection (INV-Group 5) covered 13% of the region. 20% to 34% of their areas were covered by multi-species highly suitable habitat (24 to 57% for suitable area). 11%, 21% and 1% are covered by highly suitable habitat of *T. marmoratus, R. temporaria* and *P. Punctatus* respectively (Table 4). INV-Group 5 covered the most suitable and highly suitable habitat of all studied groups and species except for *P. punctatus* with a poor overlapping between this species issues and areas of INV-Group 5 (i.e., 2%).

### 3.4 Green infrastructure (GI)

GI included and covered a larger area of highly suitable and suitable habitat for amphibian species than PA particularly hedgerow networks BOCAGE followed by WOOD and WET (see Table 2). Globally, GI covered around 50% of potential habitat for amphibian species. Among the five GI’s classes (Table 4), GI relating to traditional hedgerow landscapes (BOCAGE) were the most widespread and covered from 16% to 33% of highly suitable habitat of species or species groups (especially *T. marmoratus* with 33% and excluding *P. punctatus* with only 2%). In addition, the GI BOCAGE contained 42% highly suitable habitat for generalist species and between 12-21% for other species except *P. punctatus*. GI relating to woodlands (WOOD) and wetlands (WET) covered more than 10% of highly suitable habitat except for *T. marmoratus* and *P. punctatus*. GI relating to coastland (COAST) and natural open grasslands (OPEN) covered small areas compared with other GI and a low area of suitable habitat for studied species.

#### Potentialmulti-habitatcontinuitiespatterns

The distribution patterns associated with highly suitable and suitable habitats are shown Fig. 1 and supplemented in Fig. 3 by a more detailed analysis of the spatial organisation and potential links between habitat patches. The central and the southern parts of the region have a few patches of highly suitable habitat, especially for forest species. Generalist species have a more homogeneous distribution of suitable habitats spreads over the region. Suitable and highly suitable habitat distribution patterns associated with forest species are mostly common with other species, except for large suitable forest patches especially at the north of the region. Indeed, 92% and 81% of highly and suitable habitat of FOR species (i.e., suitability index > 65%) are include in GEN and ALL species’ high and suitable habitat respectively.

**Fig. 3.**
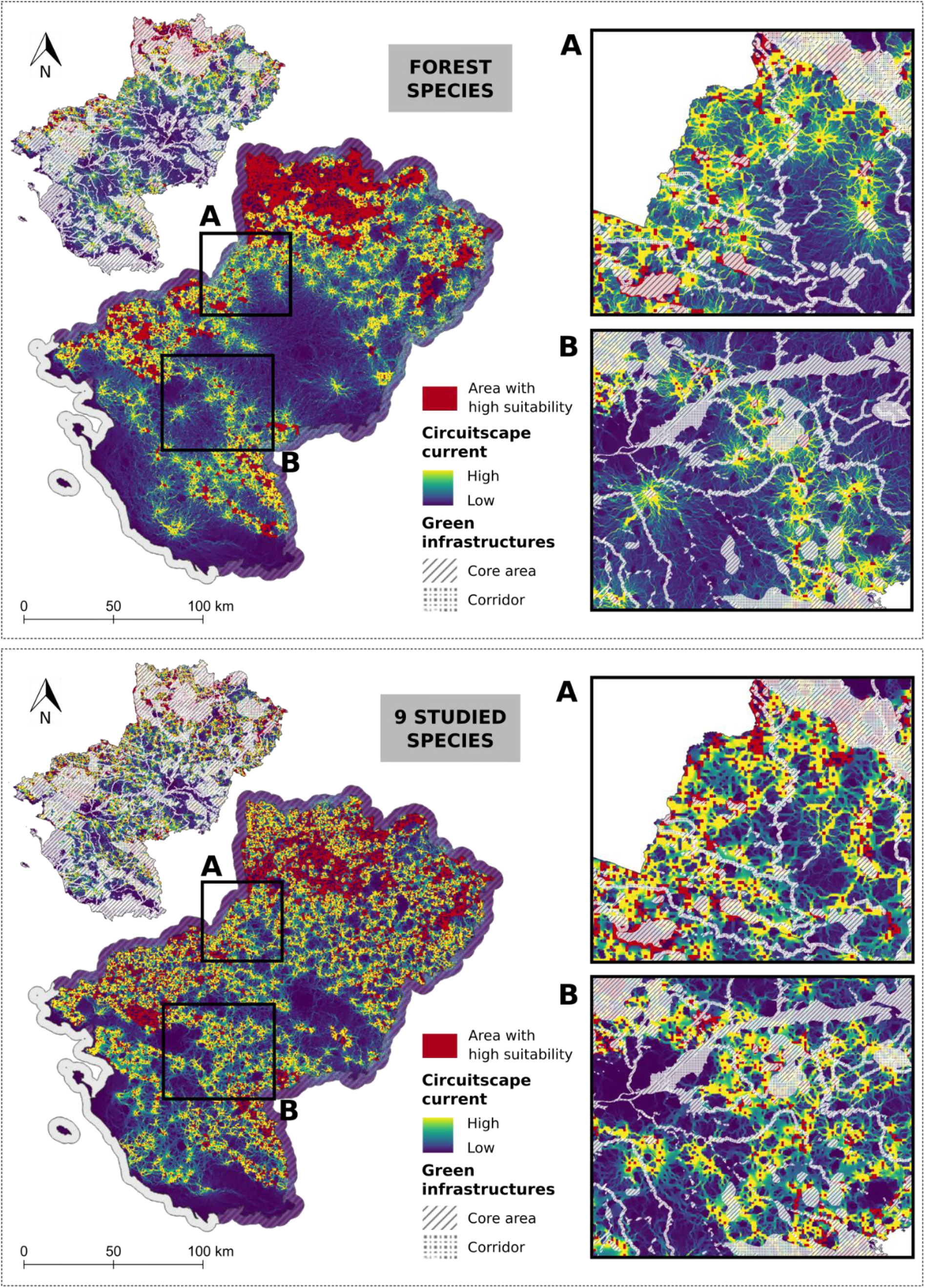
Large scale permeability between high suitability areas and associated continuities patterns confronted with regional green infrastructure. A and B show potential conflict area and issues for continuity conservation at regional scale with a mismatch with the green infrastructure network.

Regarding potential multi-specific regional continuities, two major sectors form “bottlenecks” to north-south (sector B, Fig. 3) or east-west continuity axis and potential movements (sector A, Fig. 3). This is particularly marked for forest species. These two sectors contain few protected areas and show a cover mismatch with GI (Fig. 3).

## 4 Discussion

We highlighted a mismatch between the most suitable habitats for studied species at regional scale and the existing conservation network, in particular protected areas (in coverage and efficiency). Indeed, habitat patches with a high suitability index associated with common or slightly rarer species were poorly covered by PA except for one species of high conservation concern: *P. punctatus*. Analysis of other rarer species distributions suggest that the observed mismatch concerns other species present in the studied region (Appendix 1). However, most suitable areas were better covered by GI (especially GI layers associated with wood, hedgerow landscapes and wetlands) and by INV sites (inventoried sites with ecological value but no regulatory protection). Difference in coverage and efficiency obtained for PAs and INV highlights that some sites with amphibian conservation issues are known but not protected. In addition, we identified two narrow “bottlenecks” in suitable habitat continuities at regional scale which are only very sparsely covered by the conservation network (PAs and GI).

### 4.1 General limits and interpretation framework

In addition to the limits related to the presence-only SDM detailed in Matutini et al., (2021a) recently developed stacking-SDM methods require further development to improve their evaluation. SDM are useful tools to predict the distribution of habitat suitability but rarely predict the biology of populations (Lee-Yaw et al., 2022). The multi-species networks identified should be considered simply as maps of highly suitable habitat (habitat perspective) – measured with highly sensitive and biphasic amphibian species – rather than as a real multilayer network of amphibian communities (species perspective). In addition, resistance values based on literature and expert opinions or derived from SDM values are still debated (see Godet & Clauzel, 2020 for methods comparison and discussion) and information about species movement and dispersion processes are still very scarce in the literature (Pittman et al. 2014).

Amphibians also have poor movement capacities, rarely exceeding a few kilometres during dispersal movements (Sinsch, 2014). In this work, continuities were identified from habitat distributions at regional scale and species level reflecting long-term ecological processes and population movements across decades. This study takes a first step towards assessing potential barriers to species distribution shifts in response to global change, but does not directly include landscape dynamics and climate change scenarios. Rather it provides elements at regional scale to guide regional policies and further studies to improve multi-scale integration and efficiency of measures for biodiversity conservation. Collect and integrate independent genetic data to assess our final maps, particularly in the two bottleneck sectors, would be important to evaluate connectivity and complete the investigation in view to propose relevant conservation or restoration measures.

### 4.2 A mismatch for the conservation of amphibians

Two main reasons may explain the mismatch between location of PAs and amphibian habitat requirements. Firstly, PAs are generally defined according to the needs of rare and/or emblematic species or habitats). In the study region, most PA focus on birds or plants species, large open wetlands near the western coast or the Loire River valley but only a few target species of amphibian and these tend to be rare species (e.g., *P. cultripes*, see Appendix 1). Most local, rare species of high conservation concern are pioneer species and/or associated with loose or sandy soils with poor vegetation that are found in particular on the coast, along the Loire River and in some quarries. Their ecology contrasts with the other studied species, except for *P. punctatus* which shares many of the same ecological requirements, explaining the better conservation coverage and efficiency of PA for this species.

Secondly, PA definition in the region presents a “location bias”. Indeed, PAs are defined primarily in areas of low economic interest, less used for human activities (Rodrigues et al., 2004). This bias is common worldwide and widely described in the literature (Ahmadi et al., 2020; Joppa & Pfaff, 2009). It can be especially problematic for species living in human-dominated landscapes, like most amphibian species. For example, Rodrigues et al. (2004) demonstrate that amphibians contain a large number of “gap species” worldwide, i.e., they are the biological group least covered by the global PA network. In addition, our results show a potential “network bias” similar to Ahmadi et al. (2020), i.e., the extension of PA is established without sufficiently considering their position within the network. Indeed, the current PA sites (and the sites intended for the future extension of Pas – see Appendix 5) are not located in the two areas with highest potential defined by our primary analysis of the regional continuity of suitable habitats.

### 4.3 Green infrastructure (GI) as a complementary tool for biodiversity conservation

Our results show GI offers better coverage of highly suitable habitats than PAs for amphibian species groups, but this remains partial, in particular in the two identified “bottleneck” sectors. Indeed, agricultural areas are poorly covered by PAs, generally due to a conflict of economic interest (Rodrigues et al. 2004) while GI provides complementary coverage in areas dominated by human activities. Agricultural landscapes with high conservation value for biodiversity such as hedgerow mosaic landscapes depend on the maintenance of traditional farming practices that may even be inhibited by PAs conservation (Schmitz et al., 2017).

In addition, we underline the importance of GI layers associated with wetlands, woodlands, and traditional hedgerow mosaic landscapes for amphibians. Consequently, our results highlight the importance of considering several GI layers with potential interactions. GI tends to compartmentalize conservation issues by splitting them into distinct layers of habitat types considered as independent “subnetworks”, where the “habitat” concept is limited to landcover categories (see Pilosof et al., 2017 for general perspective on multilayer nature of ecological networks). Species survival is however conditioned by different ecological processes which generally go beyond a single type of land cover class. Amphibian habitat network studies supporting GI definition mainly focus on pond networks but natural or semi-natural forest patches, especially mature forest (e.g. >70 years) and even small woodlots (Boissinot et al., 2015), are essential for amphibian population persistence, in particular in agricultural regions (Hartel et al., 2010; Collins & Fahrig 2017). Species other than amphibians may also have very contrasting requirements during their life cycle (e.g., many holometabolous insects, odonates, and bats).

Lastly, we expected a higher overlap of the GI layer called Bocage (traditional hedgerow mosaics in agricultural) based on knowledge of amphibian ecology (Boissinot et al., 2019). This GI layer was mapped using combined land use data (i.e., hedges, ponds and meadows only) at 1km-resolution with a poor consideration of the interactions between these elements (i.e., simple metrics as density or area related to hedges, ponds and meadows were combined independently). Of course, other elements of the landscape are important for amphibians, such as small streams and wooded patches, as well as more complex processes related to landscape complementation and structural heterogeneity (Boissinot et al., 2019; Collins & Fahrig 2017).

Therefore, an important weakness of GI is its exclusive reliance on land use data without including biological data and landscape complementation as well as the multilayered nature of ecological networks. Ecological niche modelling tools such as SDM, coupled with ecological connectivity models, are of particular interest for improving the ecological realism of identified networks and supporting related decisions (Clauzel & Godet 2020). These modelling tools need to be used and interpreted with care (e.g., within the framework specified in 4.1.).

### 4.4 Implications for conservation

First, we identified “bottleneck” sectors not covered by any conservation tools such as PA or GI despite high potential conservation value. These areas might be especially important for amphibian populations in a context of global change requiring distribution shifts to maintain suitable conditions for population viability. Indeed, according to Préau et al., (2019), important distribution shifts at regional scale are expected for certain amphibian species based on global change predictions. These two “bottlenecks” are located in a context of open fields and intensive farming which are considered by nature protection organisations to have limited potential for biodiversity. In these areas, some semi-natural elements such as small woods associated with wetlands may nonetheless play a key role in maintaining functionality and connectivity of the network at regional scale and should be strengthened. In these two areas, habitat continuities are maintained by riparian woods (sometime very thin and degraded) and associated river landscapes with a higher proportion of pastures and hedgerows, as well as small wood patches close to wetlands including ponds. Considering the general limits of this studies (see 4.1), these two areas need further investigation -especially connectivity evaluation by integrating genetic data – in order to defined appropriate conservation and restoration measures.

Overall, the protected areas network needs to be strengthened and supplemented by conservation tools more adapted to landscapes with strong interactions between biodiversity and human activities. Green infrastructure (GI) has better efficiency and coverage in particular because it concerns landscapes where natural and semi-natural habitats and human activity are closely intertwined. However, GI is not regulatory. Other effective area-based conservation measures (OECMs) are new conservation tools intended to complement protected areas (PAs) and might strengthen GI at local scale by sharing the common objective of restoring large scale connectivity (Convention on Biological Diversity, 2018). OECMs might be the complementary regulatory brick to PAs and GI with, in a European context, a better consideration and recognition of management practices and certain agroecological farming practices that are more biodiversity-friendly.

The different maps obtained from this study provide support for prioritizing certain regional policies with different objectives: (1) strengthening PA network, GI and future OECM (Appendix 6 provides more specific information to prioritize new protected or conservation area from a regional preselected set of sites according to our results); (2) identifying potential conflict points with transport networks (i.e., roads and railways); (3) spatially prioritizing actions supporting the agroecological transition, particularly in the two bottleneck areas considered as “degraded” for biodiversity. Conservation actions for amphibian conservation often focus on ponds and pond networks. The complementarity of ponds (or wetlands) and forest patches should be considered to define conservation actions (Cushman, 2006; Denoël & Lehmann, 2006; Pope et al., 2000) as well as certain associated extensive and traditional farming practices in western Europe (Boissinot et al., 2019; Hartel et al., 2010). In our region, amphibians have interactions with traditional agricultural practices and with related landscape dynamics. As multi-habitat species, amphibians have potential as biodiversity indicators of ecological quality of hedgerow network landscapes, composed of a mosaic of trees, grasslands, and ponds upon which they depend. The success of conservation measures depends on our ability to integrate biodiversity issues into farming practices for the benefit of amphibians but also many other forms of biodiversity benefiting from such landscape mosaics.

## Acknowledgements

This work would not have been possible without the support of the Pays de la Loire Herpetological Group, the CPIE Regional Union and the French BirdLife partner (LPO). We are especially grateful to Morgane Sineau and Benoit Marchadour who coordinate regional naturalists’ databases. We also acknowledge the many naturalists involved, for access to data and for additional fieldwork, especially Dorian Angot, Baptiste Gaboriau and Martin Bonhomme. We thank Jean Secondi and Aurélien Besnard for providing helpful discussion. Our work was supported by funding from L’Ecole Supérieure des Agricultures d’Angers, Angers Loire Metropole, The French Society for Ecology and Evolution (SFE^2^) and “Humanité et Biodiversité”.

## Authors’ contributions

**Florence Matutini**: Conceptualization (Equal), Data curation (Lead), Formal analysis (Lead), Funding acquisition (Supporting), Methodology (Lead), Investigation (Lead), Visualization (Lead), Writing-original draft (Lead), Writing-review & editing (Lead); **Marie-Josée Fortin**: Conceptualization (Equal), Methodology (Supporting), Supervision (Supporting), Writing-original draft (Supporting), Writing-review & editing (Supporting); **Jacques Baudry, Guillaume Pain & Josephine Pithon**: Conceptualization (Equal), Supervision (Equal), Writing-original draft (Supporting), Writing-review & editing (Supporting), Funding acquisition (Lead).

## Data and codes

Data sample, scripts and maps are available online: https://doi.org/10.5281/zenodo.7096821

## Conflict of interest disclosure

The authors of this preprint declare that they have no financial conflict of interest with the content of this article.

## Notes

### Competing Interest Statement

The authors have declared no competing interest.

### Summary of Updates

Addition of PCI Ecology recommendation and badge

## References

Ahmadi, M., Farhadinia, M. S., Cushman, S. A., Hemami, M. R., Nezami Balouchi, B., Jowkar, H., & Macdonald, D. W. (2020). Species and space: a combined gap analysis to guide management planning of conservation areas. Landscape Ecology, 35(7), 1505–1517. https://doi.org/10.1007/s10980-020-01033-5

Baudry, J., Bunce, R. G. H., & Burel, F. (2000). Hedgerow diversity: An international perspective on their origin, function and management. March. https://doi.org/10.1006/jema.2000.0358

Bazin, P., & Schmutz, T. (1994). La mise en place de nos bocages en Europe et leur déclin. Revue Forestière Française, S, 115. https://doi.org/10.4267/2042/26606

Boissinot, A., Besnard, A., & Lourdais, O. (2019). Amphibian diversity in farmlands: Combined influences of breeding-site and landscape attributes in western France. Agriculture, Ecosystems & Environment, 269(September 2018), 51–61. https://doi.org/10.1016/j.agee.2018.09.016

Boissinot, A., Grillet, P., Besnard, A., & Lourdais, O. (2015). Small woods positively influence the occurrence and abundance of the common frog (Rana temporaria) in a traditional farming landscape. Amphibia Reptilia, 36(4), 417–424. https://doi.org/10.1163/15685381-00003013

Burel, F., & Baudry, J. (1995). Social, aesthetic and ecological aspects of hedgerows in rural landscapes as a framework for greenways. Landscape and Urban Planning, 33(1–3), 327–340. https://doi.org/10.1016/0169-2046(94)02026-C

Calabrese, J. M., Certain, G., Kraan, C., & Dormann, C. F. (2014). Stacking species distribution models and adjusting bias by linking them to macroecological models. Global Ecology and Biogeography, 23(1), 99–112. https://doi.org/10.1111/geb.12102

Chatzimentor, A., Apostolopoulou, E., & Mazaris, A. D. (2020). A review of green infrastructure research in Europe: Challenges and opportunities. Landscape and Urban Planning, 198(March), 103775. https://doi.org/10.1016/j.landurbplan.2020.103775

Clauzel, C., & Godet, C. (2020). Combining spatial modeling tools and biological data for improved multispecies assessment in restoration areas. Biological Conservation, 250(August), 108713. https://doi.org/10.1016/j.biocon.2020.108713

Collins, J. P., & Storfer, A. (2003). Global amphibian declines: Sorting the hypotheses. Diversity and Distributions, 9(2), 89–98. https://doi.org/10.1046/j.1472-4642.2003.00012.x

Collins, S. J., & Fahrig, L. (2017). Responses of anurans to composition and configuration of agricultural landscapes. Agriculture, Ecosystems and Environment, 239, 399–409. https://doi.org/10.1016/j.agee.2016.12.038

Cushman, S. A. (2006). Effects of habitat loss and fragmentation on amphibians: A review and prospectus. Biological Conservation, 128(2), 231–240. https://doi.org/10.1016/j.biocon.2005.09.031

De Solla, S. R., Shirose, L. J., Fernie, K. J., Barrett, G. C., Brousseau, C. S., & Bishop, C. a. (2005). Effect of sampling effort and species detectability on volunteer based anuran monitoring programs. Biological Conservation, 121, 585–594. https://doi.org/10.1016/j.biocon.2004.06.018

Denoël, M., & Lehmann, A. (2006). Multi-scale effect of landscape processes and habitat quality on newt abundance: Implications for conservation. Biological Conservation, 130(4), 495–504. https://doi.org/10.1016/j.biocon.2006.01.009

Díaz-García, J. M., Pineda, E., López-Barrera, F., & Moreno, C. E. (2017). Amphibian species and functional diversity as indicators of restoration success in tropical montane forest. Biodiversity and Conservation, 26(11), 2569–2589. https://doi.org/10.1007/s10531-017-1372-2

Dirzo, R., Young, H. S., Galetti, M., Ceballos, G., Isaac, N. J. B., & Collen, B. (2014). Defaunation in the Anthropocene. Science, 345(6195), 401–406. https://doi.org/10.1126/science.1251817

Duflot, R., Avon, C., Roche, P., & Bergès, L. (2018). Combining habitat suitability models and spatial graphs for more effective landscape conservation planning: An applied methodological framework and a species case study. Journal for Nature Conservation, 46(May 2017), 38–47. https://doi.org/10.1016/j.jnc.2018.08.005

Ferrier, S., & Guisan, A. (2006). Spatial modelling of biodiversity at the community level. Journal of Applied Ecology, 43(3), 393–404. https://doi.org/10.1111/j.1365-2664.2006.01149.x

Foltête, J. C., Savary, P., Clauzel, C., Bourgeois, M., Girardet, X., Saharoui, Y., Vuidel, G., & Garnier, S. (2020). Coupling landscape graph modeling and biological data: a review. Landscape Ecology, 7. https://doi.org/10.1007/s10980-020-00998-7

Gaston, K. J., & Fuller, R. A. (2008). Commonness, population depletion and conservation biology. Trends in Ecology and Evolution, 23(1), 14–19. https://doi.org/10.1016/j.tree.2007.11.001

Godet, C., & Clauzel, C. (2020). Comparison of landscape graph modelling methods for analysing pond network connectivity. Landscape Ecology, 9. https://doi.org/10.1007/s10980-020-01164-9

Guisan, A., & Zimmermann, N. E. (2000). Predictive habitat distribution models in ecology. Ecological Modelling, 135(2–3), 147–186. https://doi.org/10.1016/S0304-3800(00)00354-9

Hamer, A. J., & McDonnell, M. J. (2008). Amphibian ecology and conservation in the urbanising world: A review. Biological Conservation, 141(10), 2432–2449. https://doi.org/10.1016/j.biocon.2008.07.020

Hartel, T., Schweiger, O., Öllerer, K., Cogâlniceanu, D., & Arntzen, J. W. (2010). Amphibian distribution in a traditionally managed rural landscape of Eastern Europe: Probing the effect of landscape composition. Biological Conservation, 143(5), 1118–1124. https://doi.org/10.1016/j.biocon.2010.02.006

Jennings, M. D. (2000). Gap analysis: Concepts, methods, and recent results. Landscape Ecology, 15(1), 5–20. https://doi.org/10.1023/A:1008184408300

Joppa, L. N., & Pfaff, A. (2009). High and far: Biases in the location of protected areas. PLoS ONE, 4(12), 1–6. https://doi.org/10.1371/journal.pone.0008273

Keeley, A. T. H., Beier, P., & Gagnon, J. W. (2016). Estimating landscape resistance from habitat suitability: effects of data source and nonlinearities. Landscape Ecology. https://doi.org/10.1007/s10980-016-0387-5

Keeley, A. T. H., Beier, P., Keeley, B. W., & Fagan, M. E. (2017). Habitat suitability is a poor proxy for landscape connectivity during dispersal and mating movements. Landscape and Urban Planning, 161, 90–102. https://doi.org/10.1016/j.landurbplan.2017.01.007

Kleijn, D., Rundlöf, M., Scheper, J., Smith, H. G., & Tscharntke, T. (2011). Does conservation on farmland contribute to halting the biodiversity decline? Trends in Ecology and Evolution, 26(9), 474–481. https://doi.org/10.1016/j.tree.2011.05.009

Koen, E. L., Bowman, J., Sadowski, C., & Walpole, A. A. (2014). Landscape connectivity for wildlife: Development and validation of multispecies linkage maps. Methods in Ecology and Evolution, 5(7), 626–633. https://doi.org/10.1111/2041-210X.12197

Lee-Yaw, J. A., McCune, J. L., Pironon, S., & Sheth, S. N. (2022). Species distribution models rarely predict the biology of real populations. Ecography, 2022(6), 1–16. https://doi.org/10.1111/ecog.05877

Matutini, F., Baudry, J., Fortin, M. J., Pain, G., & Pithon, J. (2021a). Integrating landscape resistance and multi-scale predictor of habitat selection for amphibian distribution modelling at large scale. Landscape Ecology, 36(12), 3557–3573. https://doi.org/10.1007/s10980-021-01327-2

Matutini, F., Baudry, J., Pain, G., Sineau, M., & Pithon, J. (2021b). How citizen science could improve species distribution models and their independent assessment. Ecology and Evolution, November 2020, ece3.7210. https://doi.org/10.1002/ece3.7210

Mazerolle, M. J. (2005). Drainage ditches facilitate frog movements in a hostile landscape. Landscape Ecology, 20(5), 579–590. https://doi.org/10.1007/s10980-004-3977-6

McRae, B. H.,., Dickson, B. G., Keitt, T. H., & Shah, V. B. (2008). Using Circuit Theory to Model Connectivity in Ecology, Evolution, and Conservation. Ecology, 89(10), 2712–2724.

McRae, B., Shah, V., & Edelman, A. (2016). Circuitscape: modeling landscape connectivity to promote conservation and human health. The Nature Conservancy, 1–14.

Moilanen, A., Franco, A. M. A., Early, R. I., Fox, R., Wintle, B., & Thomas, C. D. (2005). Prioritizing multiple-use landscapes for conservation: Methods for large multi-species planning problems. Proceedings of the Royal Society B: Biological Sciences, 272(1575), 1885–1891. https://doi.org/10.1098/rspb.2005.3164

Opdam, P., Steingröver, E., & Rooij, S. Van. (2006). Ecological networks: A spatial concept for multi-actor planning of sustainable landscapes. Landscape and Urban Planning, 75(3–4), 322–332. https://doi.org/10.1016/j.landurbplan.2005.02.015

Petrovan, S. O., & Schmidt, B. R. (2016). Volunteer conservation action data reveals large-scale and long-term negative population trends of a widespread amphibian, the common toad (Bufo bufo). PLoS ONE, 11(10), 1–12. https://doi.org/10.1371/journal.pone.0161943

Pilosof, S., Porter, M. A., Pascual, M., & Kéfi, S. (2017). The multilayer nature of ecological networks. In Nature Ecology and Evolution (Vol. 1, Issue 4). Nature Publishing Group. https://doi.org/10.1038/s41559-017-0101

Pope, S. E., Fahrig, L., & Merriam, H. G. (2000). Landscape complementation and metapopulation effects on leopard frog populations. Ecology, 81(9), 2498–2508. https://doi.org/10.1890/0012-9658(2000)081[2498:LCAMEO]2.0.CO;2

Rodrigues, A. S. L., & Cazalis, V. (2020). The multifaceted challenge of evaluating protected area effectiveness. Nature Communications, 1–4. https://doi.org/10.1038/s41467-020-18989-2

Rodrigues, A. S., Andelman, S. J., Bakarr, M. I., Boitani, L., Brooks, T. M., Cowling, R. M., … & Yan, X. (2004). Effectiveness of the global protected area network in representing species diversity. Nature, 428(6983), 640–643. https://doi.org/10.1038/nature02459.1.

Salomaa, A., Paloniemi, R., Kotiaho, J. S., Kettunen, M., Apostolopoulou, E., & Cent, J. (2017). Can green infrastructure help to conserve biodiversity? Environment and Planning C: Government and Policy, 35(2), 265–288. https://doi.org/10.1177/0263774X16649363

Scherrer, D., D’Amen, M., Fernandes, R. F., Mateo, R. G., & Guisan, A. (2018). How to best threshold and validate stacked species assemblages? Community optimisation might hold the answer. Methods in Ecology and Evolution, 9(10), 2155–2166. https://doi.org/10.1111/2041-210X.13041

Scherrer, D., Mod, H. K., & Guisan, A. (2020). How to evaluate community predictions without thresholding? Methods in Ecology and Evolution, 11(1), 51–63. https://doi.org/10.1111/2041-210X.13312

Schmitz, M. F., Herrero-Jáuregui, C., Arnaiz-Schmitz, C., Sánchez, I. A., Rescia, A. J., & Pineda, F. D. (2017). Evaluating the Role of a Protected Area on Hedgerow Conservation: The Case of a Spanish Cultural Landscape. Land Degradation and Development, 28(3), 833–842. https://doi.org/10.1002/ldr.2659

Seibold, S., Gossner, M. M., Simons, N. K., Blüthgen, N., Müller, J., Ambarli, D., Ammer, C., Bauhus, J., Fischer, M., Habel, J. C., Linsenmair, K. E., Nauss, T., Penone, C., Prati, D., Schall, P., Schulze, E. D., Vogt, J., Wöllauer, S., & Weisser, W. W. (2019). Arthropod decline in grasslands and forests is associated with landscape-level drivers. Nature, 574(7780), 671–674. https://doi.org/10.1038/s41586-019-1684-3

Sewell, D., & Griffiths, R. A. (2009). Can a single amphibian species be a good biodiversity indicator? Diversity, 1(2), 102–117. https://doi.org/10.3390/d1020102

Sinsch, U. (2014). Movement ecology of amphibians: from individual migratory behaviour to spatially structured populations in heterogeneous landscapes 1, 2. Canadian Journal of Zoology, 92(6), 491–502. https://doi.org/10.1139/cjz-2013-0028

Snäll, T., Lehtomäki, J., Arponen, A., Elith, J., & Moilanen, A. (2016). Green Infrastructure Design Based on Spatial Conservation Prioritization and Modeling of Biodiversity Features and Ecosystem Services. Environmental Management, 57(2), 251–256. https://doi.org/10.1007/s00267-015-0613-y

Stuart, S. N., Chanson, J. S., Cox, N. A., Young, B. E., Rodrigues, A. S. L., Fischman, D. L., & Waller, R. W. (2004). Status and trends of amphibian declines and extinctions worldwide. Science, 306(5702), 1783–1786. https://doi.org/10.1126/science.1103538

Thuiller, W., Pollock, L. J., Gueguen, M., & Münkemüller, T. (2015). From species distributions to meta-communities. 18(12), 1321–1328. https://doi.org/10.1111/ele.12526.

