## Appendix 1 for "Conservation networks do not match the ecological requirements of amphibians"

**Appendix 1. species data, supplementary analysis on other species and model transferability**

1. **Data used for SDM calibration and validation (completed from Matutini et al. 2021)**

*Opportunistic presence data (calibration dataset)*

**Table S1 Data sources for model calibration**

| **Database name** | **Type** | **General website** | **Data proportion** |
| --- | --- | --- | --- |
| Faune_anjou | Citizen bases with validation process by professionals | <https://www.faune-anjou.org/> | 25% |
| Faune_maine |  | https://www.faune-maine.org/ | 10% |
| Faune_vendee |  | <https://www.faune-vendee.org/> | 10% |
| Faune_loire_atlantique |  | <https://www.faune-loire-atlantique.org/> | 11% |
| Biolovision |  | https://data.biolovision.net/ | 18% |
| URCPIE | Professional & volunteers | <http://urcpie-paysdelaloire.org/> | 12% |
| Bretagne Vivante | Naturalist group | <https://www.bretagne-vivante.org/> | 2% |
| ONF_BDN | Professional | <https://www.onf.fr/> | 3% |
| SICEN | Professional | <http://www.cenpaysdelaloire.fr/> | 3% |
| BASEPARC PNRMP / OPN | Professional | <https://pnr.parc-marais-poitevin.fr/> | 2% |
| Naturalistes en lutte | Naturalist group | <https://naturalistesenlutte.wordpress.com/> | 2% |
| Sterne 2.0 | Professional | <http://www.sterne2.com/> | 1% |
| Les naturalistes vendeens | Naturalist group | <http://naturalistes-vendeens.org/> | 1% |
| Gouret_FLA | Naturalist individual base | **-** | <1% |
| Cap Atlantique | Professional | <https://www.cap-atlantique.fr/accueil> | <1% |
| Undragon.org | Citizen base | <http://undragon.org/> | <1% |
| ONCFS | Professional | <http://www.oncfs.gouv.fr/> | <1% |

**Table S2. Description of the presence-only data used for each of nine species for calibration of habitat suitability models**.

| **Species** | Opportunistic presence-only dataset  (model calibration and cross- validation) | | |
| --- | --- | --- | --- |
|  | Total nb of presence | | Nb of 500 m presence-cells |
| **Anurans:** |  | |  |
| *Bufo spinosus* | 8320 | | 4127 |
| *Hyla arborea arborea* | 6344 | | 3353 |
| *Pelodytes punctatus* | 2711 | | 1103 |
| *Rana dalmatina* | 9073 | | 3752 |
| *Rana temporaria* | 1525 | | 477 |
| **Urodeles:** |  |  |  |
| *Salamandra Salamandra terrestris* | 4916 | | 2242 |
| *Triturus marmoratus* | 1478 | | 629 |
| *Triturus cristatus* | 1791 | | 766 |
| *Lissotriton helveticus* | 7047 | | 2835 |

*Standardised detection-nondetection data (external validation dataset)*

**Name of the citizen science program**: “Un Dragon dans mon Jardin”

**Coordination**: URCPIE – “Union régionale des centres d’initiatives pour l’environnement ».

For external SDM validation, we extracted detection-nondetection amphibian data from a regional citizen science database. This database contained 576 monitored aquatic sites for the period 2013-2019, with observations made in the context of a programme aiming to estimate amphibian population trends (regionally called “Un Dragon dans mon Jardin”). Observers had to follow a standard protocol; each site had to be monitored three times separated by at least one month - one diurnal between January and March and two nocturnal between March and June – to cover different species’ breeding periods, during good weather conditions (no frost, no rain, no or weak wind). For each survey, three complementary methods were used to detect amphibians: an acoustic survey (5 min at 5 metres from the site without light) to detect breeding calls of male Anurans specie; an active visual survey using a flashlight torch (500-1000 lumens) to observe individuals and eggs and a catching survey using a net (3 net sweeps per site) if the observer had specific authorization. These methods are commonly used for amphibian community surveys. If the protocol was not respected and/or if the observer was not sufficiently experienced, only the presence data were considered valid (exclusion of the absence data). In addition, a threshold value for minimum sampling effort required were used to validate non-detection as absence data. See Matutini et al. 2020

**Table S3. Description of the filtered datasets for each of nine species used for external validation of habitat suitability models**. DET: 500 m cells with detection of the species; NoDET: 500m nondetection-cells. Results for 1 interaction

|  | EVAL | | | | EVAL_STRAT |
| --- | --- | --- | --- | --- | --- |
|  | **Nb of**  **DET** | | **Nb of**  **NoDET** | | Nb data/strat |
| **Anurans:** |  | |  | |  |
| *Bufo spinosus* | 97 | | 187 | | 23 |
| *Hyla arborea arborea* | 136 | | 204 | | 23 |
| *Pelodytes punctatus* | 40 | | 249 | | 8 |
| *Rana dalmatina* | 186 | | 162 | | 30 |
| *Rana temporaria* | 17 | | 231 | | 15 |
| **Urodeles:** |  | |  | |  |
| *Salamandra Salamandra terrestris* | 79 | | 186 | | 25 |
| *Triturus marmoratus* | 62 | | 213 | | 16 |
| *Triturus cristatus* | 52 | | 241 | | 25 |
| *Lissotriton helveticus* | 176 | | 164 | | 13 |

**Figure S1. Distribution of the 500m² cells with data for the opportunistic dataset and the external evaluation dataset (e.g. *CS.1+ABS+SUP*).** (1) All 500m² cells with at least one opportunistic observation (all species); (2) 500m² cells used as presence-absence data (with at least three surveys performed by an expert observer or six surveys by an intermediate observer) for external validation; (3) 500m² cells used only as presence if the species had been detected (sampling effort too weak for absence data) for external validation. The external dataset for validation is a compilation of (2) (presence-absence) and (3) (presence).

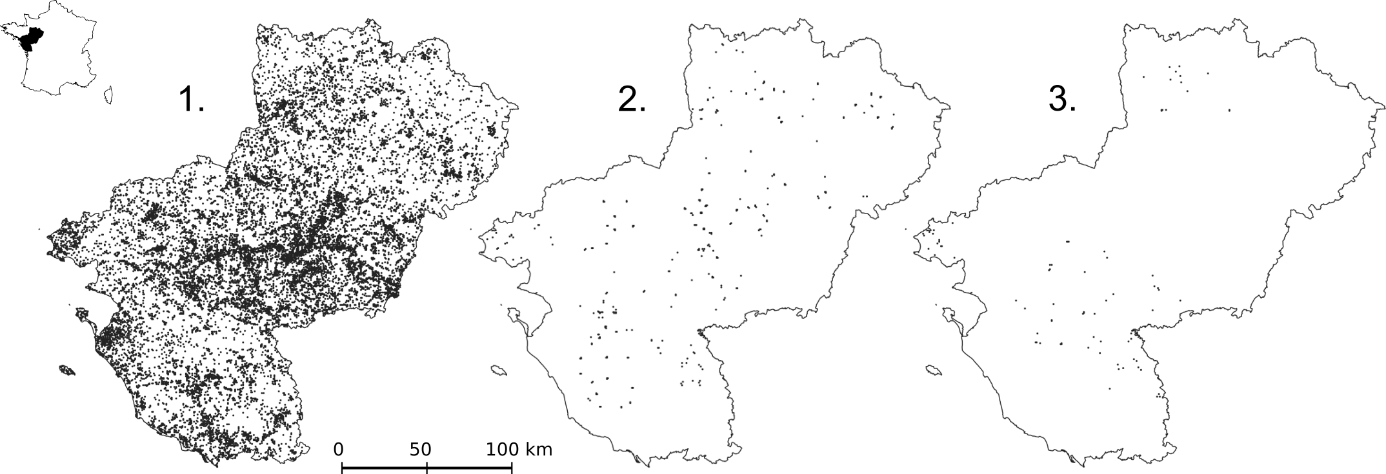

2. **Supplementary analysis on other species and model transferability**
   2.1. **Species description**

Among all the species present in Pays-de-la-Loire described below, we have excluded from this analysis:

- Exotic and/or invasive species;
- Rare species at the limit of the distribution area (i.e. with very little data at the edge of the study area) and/or species with less than 5 points;
- Species of the Pelophylax genus subject to identification errors because they are morphologically very similar and can hybridize. Given the number of observations and conservation issues, Pelophylax kl. esculentus was however included in the analysis (expert data with validated identification only).

**Table S4. Description of the amphibian’s species of Pays-de-la-Loire.** RL FR: France red list (2015); RL Region: regional red list of Pays-de-la-Loire (2021); Priority: regional priority level for species conservation from the regional red list from 0 (low) to 3 (very high) defined by Marchadour et al. (2021); Hab. Dir. N2000: habitats directive classification (Natura 2000); Nb presence data: Number of presence-only data were calculated after a 100 m-resolution rasterization and correspond to the number of pixels (cells) with at least one observation of the species. Species names with an asterisk (*) has been included to the complementary analysis (see exclusion criteria presented above).

|  | Code | Species | RL FR | RL Region | Regional priority (2009/2021) | Hab. Dir. N2000 | Nb presence data (Atlas) – 100m-cell | Nb presence data (Atlas) – 500m-cell |
| --- | --- | --- | --- | --- | --- | --- | --- | --- |
| Species used for SSDM | *PELPUN* | *Pelodytes punctatus* | LC | NT | 2/1 |  | 1828 | 1316 |
|  | ***TRIMAR*** | ***Triturus marmoratus*** | **NT** | **NT** | **3/3** | **IV** | **890** | **703** |
|  | *RANTEM* | *Rana temporaria* | LC | VU | 2/1 | V | 891 | 515 |
|  | *LISHEL* | *Lissotriton helveticus* | LC | LC | 1/1 |  | 4509 | 3279 |
|  | *BUFSPI* | *Bufo spinosus* | / | LC | 0/1 |  | 6772 | 5006 |
|  | *HYLARB* | *Hyla arborea* | NT | LC | 0/1 | IV | 5026 | 3919 |
|  | *RANDAL* | *Rana dalmatina* | LC | LC | 0/1 | IV | 6275 | 4481 |
|  | *TRICRI* | *Triturus cristatus* | NT | NT | 0/2 | II & IV | 1126 | 894 |
|  | *SALSAL* | *Salamandra salamandra* | LC | LC | 0/0 |  | 3703 | 2625 |
| Other species | *EPICAL* | *Epidalea* *calamita** | LC | NT | 2/1 | IV | 443 | 275 |
|  | ***LISVUL*** | ***Lissotriton* *vulgaris**** | **NT** | **EN** | **2/2** |  | **116** | **102** |
|  | *ICHALP* | *Ichthyosaura* *alpestris** | LC | NT | 2/0 |  | 480 | 346 |
|  | *ALYOBS* | *Alytes obstetricans** | LC | NT | 1/1 | IV | 1059 | 820 |
|  | *TRIBLA* | *Triturus x blasii** | / | / | / | / | 79 | 69 |
|  | ***/*** | ***Bombina variegata*** | **VU** | **CR** | **3/3** | **II & IV** | **31** | **13** |
|  | ***/*** | ***Pelobates* *cultripes*** | **VU** | **EN** | **3/3** | **IV** | **127** | **48** |
|  | */* | *Hyla meridionalis* | LC | LC | 1/0 | IV | 323 | 229 |
|  | */* | *Xenopus laevis* | / | / | / |  | 131 | 112 |
|  | ***/*** | ***Pelophylax lessonae*** | **NT** | **VU** | **3/2** | **IV** | **5** | **5** |
|  | ***PELESC*** | ***Pelophylax kl. esculentus**** | **NT** | **NT** | **0/2** | **V** | **63** | **60** |
|  | */* | *Pelophylax ridibundus* | LC | / | / | V | 140 | 131 |
|  | ***/*** | ***Pelophylax perezi*** | **NT** | **EN** | **0/2** | **V** | **1** | **1** |
|  | ***/*** | ***Pelophylax kl. grafi*** | **NT** | **EN** | **0/2** |  |  |  |
|  | ***/*** | ***Pelophylax sp.*** | **/** | **/** | **/** | **/** | **10390** | **7277** |

* species included in the complementary analysis Figure 2

2.2. **Transferability of SSDM to rarer species**

**Figure S1.2. Distribution of presence-only data of selected species according to the different suitability gradients modelled (SDM) for two species (PELPUN and TRIMAR) and for tree species groups (Stack-SDM for ALL, GEN and FOR).** Lines show the two thresholds 65% and 80% in the suitability index. For species code, see Table 1. PELPUN and TRIMAR are the two species considered by thresholders as model species to asses regional ecological network functionality.

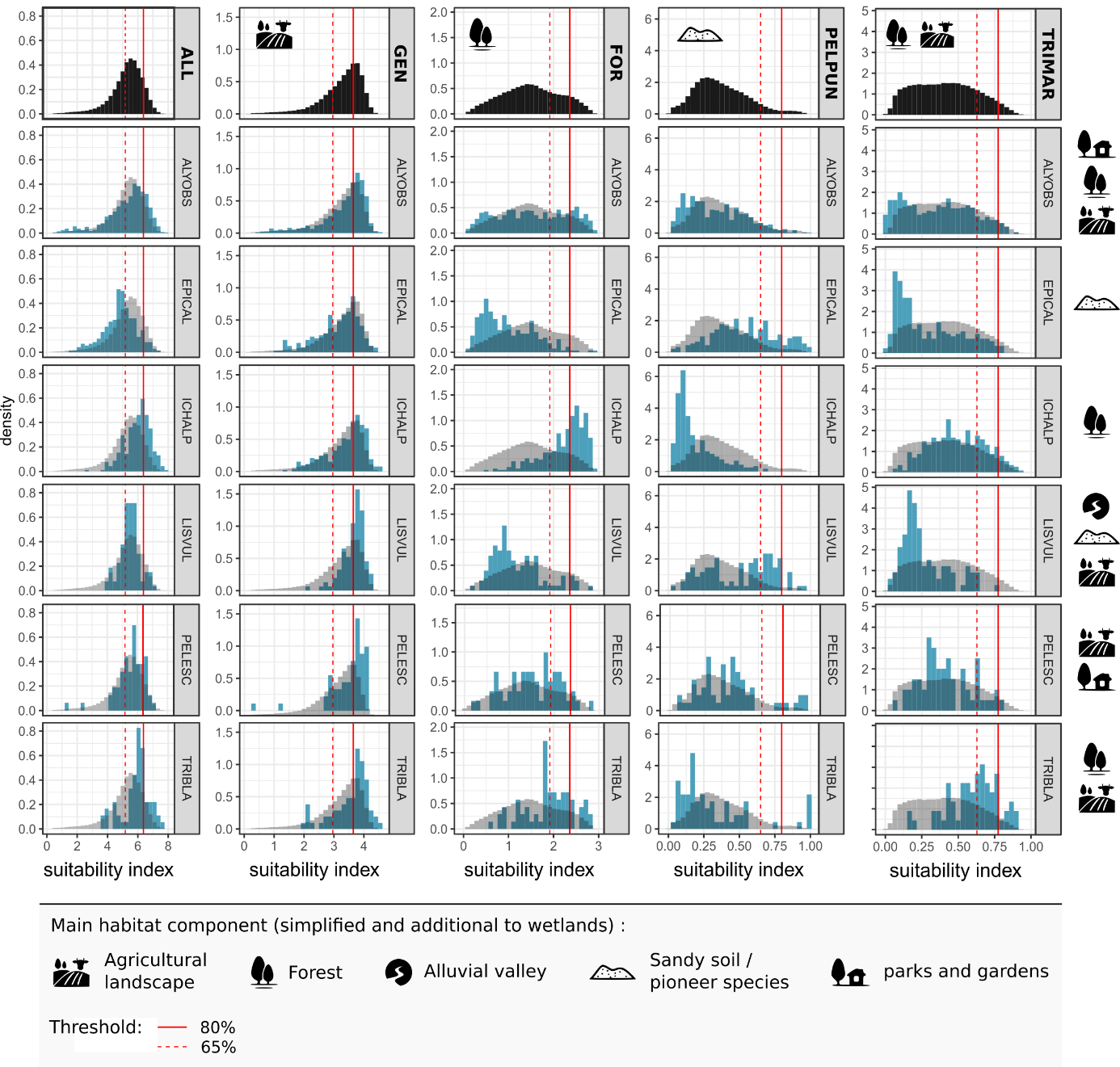

2.3. **Conservation coverage for rarer species (with presence data only)**

**Table S1.5****. Proportion of presence data covered by the conservation area (conservation coverage) for species not included in species distribution modelling.** 1. Presence data used in the analysis are defined as a 100m-pixel with at least one observation of the species. 2. Same as 1 but we add a distance condition of 500m between each species observations. See Table 2 (main text) for “code names” details and Table 1 for species names. * show species at limit of their geographic ranges or species with very few presence data.

| **1.** |  |  | **No minimal distance between 100m pixel with at least 1 presence data** | | | | | | | | |
| --- | --- | --- | --- | --- | --- | --- | --- | --- | --- | --- | --- |
| **Grp** | **Name** | **classe_UICN** | **ALYOBS** | **EPICAL** | **LISVUL** | **ICHALP** | **TRIBLA** | **PELESC** | **PELCUL*** | **BOMVAR*** | **PELLES*** |
| 1 | RNN | I_IV | 0,00 | 0,01 | 0,00 | 0,00 | 0,00 | 0,00 | 0,03 | 0,00 | 0,00 |
| 1 | SIN_SCL | III | 0,04 | 0,09 | 0,07 | 0,00 | 0,01 | 0,00 | 0,28 | 0,00 | 0,00 |
| 1 | RNR_RB_SLit_APB | IV | 0,00 | 0,03 | 0,01 | 0,01 | 0,02 | 0,00 | 0,06 | 0,00 | 0,00 |
| 2 | SCEN_PNR | V | 0,10 | 0,34 | 0,05 | 0,08 | 0,07 | 0,16 | 0,16 | 0,02 | 0,00 |
| 3 | N2000 | INT | 0,06 | 0,10 | 0,44 | 0,16 | 0,12 | 0,06 | 0,19 | 0,00 | 0,20 |
| 4 | BOCAGE | GI | 0,09 | 0,02 | 0,05 | 0,14 | 0,15 | 0,10 | 0,02 | 0,12 | 0,00 |
| 4 | WOOD | GI | 0,07 | 0,07 | 0,02 | 0,32 | 0,12 | 0,05 | 0,00 | 0,00 | 0,00 |
| 4 | WET | GI | 0,09 | 0,09 | 0,12 | 0,24 | 0,20 | 0,14 | 0,00 | 0,04 | 0,00 |
| 4 | COAST | GI | 0,00 | 0,04 | 0,00 | 0,00 | 0,00 | 0,00 | 0,00 | 0,00 | 0,00 |
| 4 | OPEN | GI | 0,01 | 0,03 | 0,02 | 0,00 | 0,01 | 0,00 | 0,00 | 0,00 | 0,00 |
| 4 | Total core hab/ | GI | 0,19 | 0,15 | 0,16 | 0,47 | 0,39 | 0,19 | 0,02 | 0,12 | 0,00 |
| 4 | Corridor | GI | 0,11 | 0,04 | 0,05 | 0,09 | 0,06 | 0,16 | 0,00 | 0,51 | 0,40 |
| 5 | ZNIEFF | INV | 0,08 | 0,11 | 0,10 | 0,31 | 0,33 | 0,06 | 0,05 | 0,04 | 0,00 |
| / | SCAP | PROP | 0,14 | 0,41 | 0,43 | 0,21 | 0,14 | 0,17 | 0,77 | 0,00 | 0,20 |
|  | **TOTAL REGION (nb data)** | | **1160** | **504** | **122** | **541** | **84** | **63** | **187** | **31** | **5** |
| **2.** |  |  | **Minimal distance of 500m between 100m pixel with at least 1 presence data** | | | | | | | | |
| **Groupe** | **PA** | **classe_UICN** | **ALYOBS** | **EPICAL** | **LISVUL** | **ICHALP** | **TRIBLA** | **PELESC** | **PELCUL*** | **BOMVAR*** | **PELLES*** |
| 1 | RNN | I_IV | 0,00 | 0,00 | 0,00 | 0,00 | 0,00 | 0,00 | 0,00 | 0,00 | 0,00 |
| 1 | SIN_SCL | III | 0,04 | 0,09 | 0,07 | 0,01 | 0,02 | 0,00 | 0,21 | 0,00 | 0,00 |
| 1 | RNR_RB_SLit_APB | IV | 0,00 | 0,02 | 0,01 | 0,01 | 0,03 | 0,00 | 0,07 | 0,00 | 0,00 |
| 2 | SCEN_PNR | V | 0,10 | 0,33 | 0,06 | 0,10 | 0,08 | 0,15 | 0,17 | 0,14 | 0,00 |
| 3 | N2000 | INT | 0,07 | 0,12 | 0,36 | 0,13 | 0,11 | 0,06 | 0,24 | 0,00 | 0,25 |
| 4 | BOCAGE | GI | 0,12 | 0,02 | 0,06 | 0,14 | 0,16 | 0,09 | 0,03 | 0,00 | 0,00 |
| 4 | WOOD | GI | 0,07 | 0,07 | 0,02 | 0,23 | 0,11 | 0,06 | 0,00 | 0,00 | 0,00 |
| 4 | WET | GI | 0,09 | 0,12 | 0,13 | 0,15 | 0,17 | 0,13 | 0,00 | 0,00 | 0,00 |
| 4 | COAST | GI | 0,00 | 0,02 | 0,00 | 0,00 | 0,00 | 0,00 | 0,00 | 0,00 | 0,00 |
| 4 | OPEN | GI | 0,01 | 0,02 | 0,02 | 0,00 | 0,02 | 0,00 | 0,00 | 0,00 | 0,00 |
| 4 | Total core hab/ | GI | 0,21 | 0,17 | 0,19 | 0,39 | 0,38 | 0,19 | 0,03 | 0,00 | 0,00 |
| 4 | Corridor | GI | 0,10 | 0,04 | 0,07 | 0,11 | 0,08 | 0,17 | 0,00 | 0,14 | 0,50 |
| 5 | ZNIEFF | INV | 0,09 | 0,13 | 0,11 | 0,25 | 0,27 | 0,06 | 0,07 | 0,14 | 0,00 |
| / | SCAP | PROP | 0,14 | 0,37 | 0,38 | 0,13 | 0,16 | 0,17 | 0,76 | 0,00 | 0,25 |
|  | **TOTAL REGION (nb data)** | | **693** | **211** | **90** | **272** | **63** | **54** | **29** | **7** | **4** |
