## Appendix 2 for "Conservation networks do not match the ecological requirements of amphibians"

**Appendix 2: studied conservation network in the region Pays-de-la-Loire**

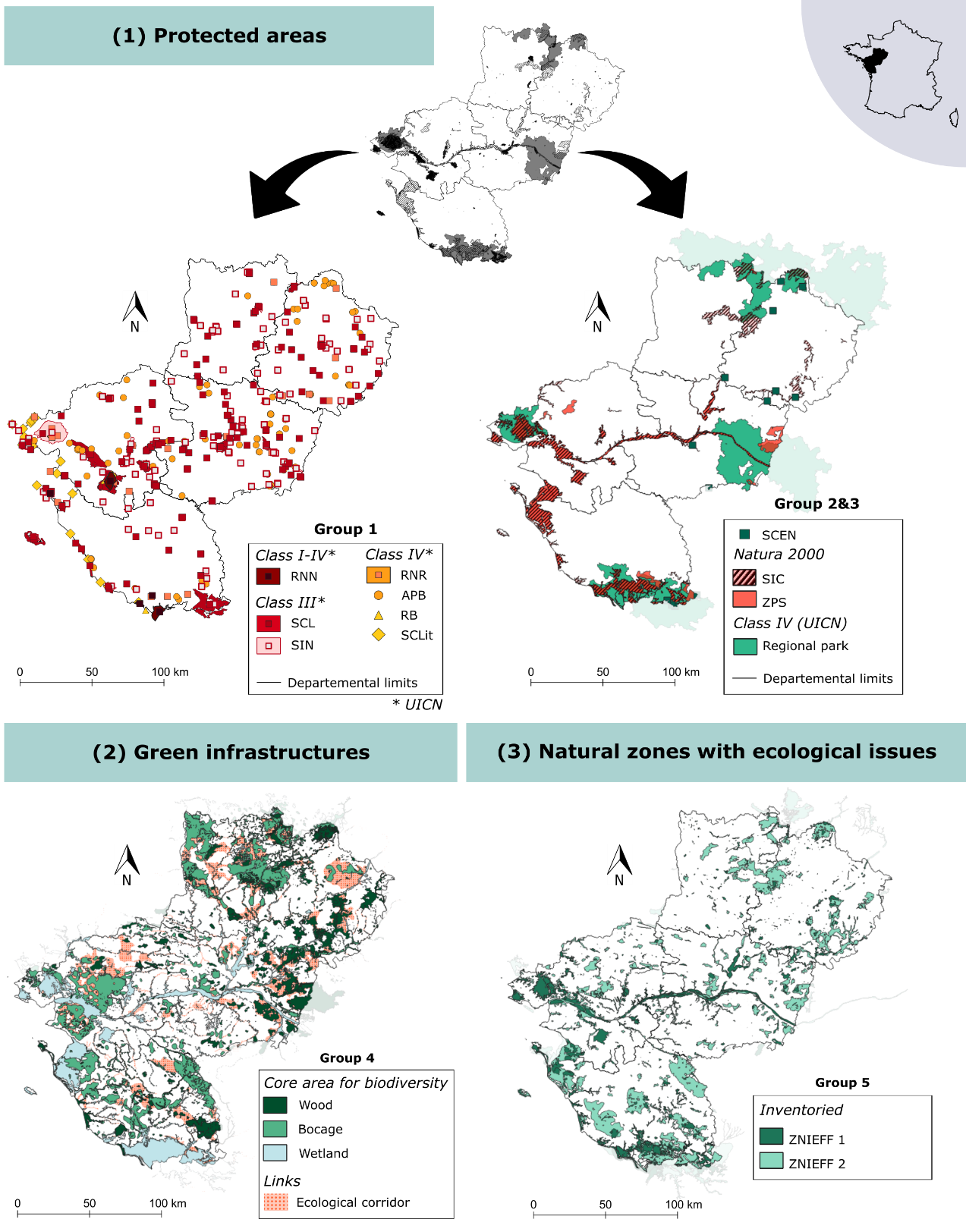

**Figure S1. Existing conservation area network including protected area (PA), Green infrastructure (GI) and surveyed natural zones with ecological value (INV).** Groups and categories are detailed Table 2. The GI named COAST and OPEN was not mapped because of their low surface and a secondary ecological interest for conservation of studied species.
