## Appendix 3 for "Conservation networks do not match the ecological requirements of amphibians"

**Appendix 3: Method selection for Stacking-SDM**

**Table S1. Accuracy of stacking species distribution models using different stacking methods. Results obtain with 100 permutations.** Bold values show selected model and coloured cells show model uses for further analysis.

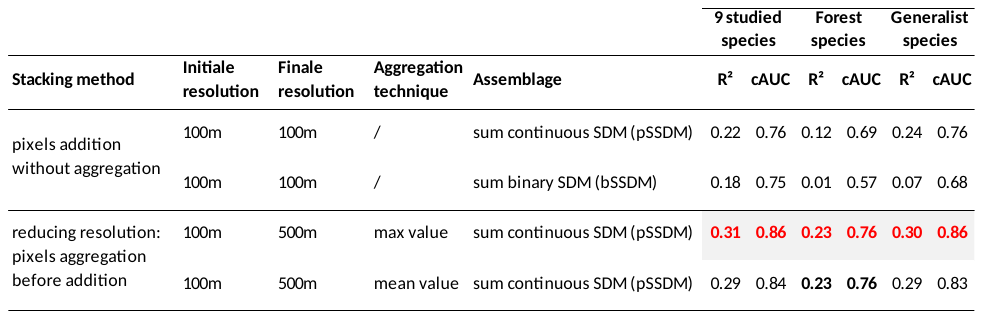
