## Appendix 4 for "Conservation networks do not match the ecological requirements of amphibians"

**Appendix 4: Friction values attribution**

**Table S1. Friction values based on literature and expert opinions**

|  | **Land cover** | **Description** | **Friction cost** |
| --- | --- | --- | --- |
| **1. Barrier without pass** | Railway for high-speed train | Pixel with a railway for high-speed train | 200 |
|  | Stream>50 m | Pixel with a river larger than 50 m | 200 |
|  | Topographic barriers (Slope>45%) | Pixel with high slope, mainly rare cliffs and gorges | 200 |
|  | Urban (high density) | Pixel with 100% of dense urban area | 200 |
|  | Primary road (highway and dual carriageways) | Pixel with a high traffic road as highway and dual carriageways | 200 |
|  | Railway | Pixel with a railway | 100 |
| **2. Linear barrier with underway pass** | Underway pass: |  |  |
|  | secondary road underpass |  | 60 |
|  | Pipe |  | 60 |
|  | Stream |  | 10 |
|  | Path/track |  | 50 |
|  | Wildlife bridges |  | 20 |
|  | Viaduc |  | 1-20 |
| **3. Presence of pond/lake**  **If not -> 4** | Pond: |  |  |
|  | +wood>50% | Pixel with at least 1 pond and more than 50% of woods | 1 |
|  | +wood20-50% | Pixel with at least 1 pond and small amount of woods | 2 |
|  | +hedgerow | Pixel with at least 1 pond and 50 m hedgerows | 5 |
|  | +meadow | Pixel with at least 1 pond, no tree but more than 50% of meadow | 10 |
|  | +culture 100% | Pixel with at least 1 pond, no tree, in a 75% crop context | 20 |
|  | +urban | Pixel with at least 1 pond, no tree, in a 100% urban context | 40 |
| **4. Presence of stream**  **If not -> 5** | Stream <50 m: |  |  |
|  | +wood | Pixel with a small stream and at least 50% of wood | 1 |
|  | +hedgerow | Pixel with a small stream and at least 50 m of hedgerows | 2 |
|  | Wood<50 and/or Hedgerow<50 and meadow | Pixel with a small stream and few trees in meadow context (>50%) | 2 |
|  | no tree (nt) + meadow | Pixel with a small stream and no tree | 5 |
|  | +nt+crop 100% | Pixel with a small stream and no tree in crop context (>50%) | 20 |
|  | +urban dense | Pixel with a small stream and no tree in urban context (>50%) | 30 |
| **5. other class (no stream and no pond)** | Path with hedgerows (ditches) | Pixel with a path with at least 1 side with hedgerow | 5 |
|  | Path without trees (ditches) + non urban | Pixel with a path in a non-urban open area | 10 |
|  | Riparian forest | Pixel with a riparian forest (75%) | 2 |
|  | Deciduous and mixed wood | Pixel with more 75% of deciduous and or mixed wood | 5 |
|  | Evergreen forest | Pixel with more 75% of evergreen wood | 5 |
|  | Meadow | Pixel with more than 50% of meadow | 15 |
|  | Meadow + wood | Pixel with more than 50% of meadow and small wood | 10 |
|  | Meadow in high MPH (humidity index) | Pixel with more than 50% of Wet meadow (MPH>=1) | 10 |
|  | Meadow in high MPH (humidity index) + hedgerow or shrubs | Wet meadow (MPH>=1) with at least 50 m hedgerow and/or shrubs | 5 |
|  | Meadow in sandy soil (>50%) | Pixel with more than 50% of meadow aver sandy soil | 20 |
|  | Shrubs | Pixel with more 75% of shrubs | 20 |
|  | Stream from 15 m to 50 m | Pixel with a stream large from 15 to 50 | 50 |
|  | Crop 100% | Pixel with 100% of crops | 50 |
|  | Crop + urban 100% | Pixel with 100% of crop/urban | 20 |
|  | Secondary roads | Pixel with a secondary road | 50 |
|  | Urban low density (no farm) | Pixel with 100% of low dense urban area (houses + gardens) | 50 |
| Un classed pixels (<5%) |  |  | 20 |
