## Appendix 5 for "Conservation networks do not match the ecological requirements of amphibians"

**Appendix 5 – Confidence maps and sensitivity to thresholds selection**

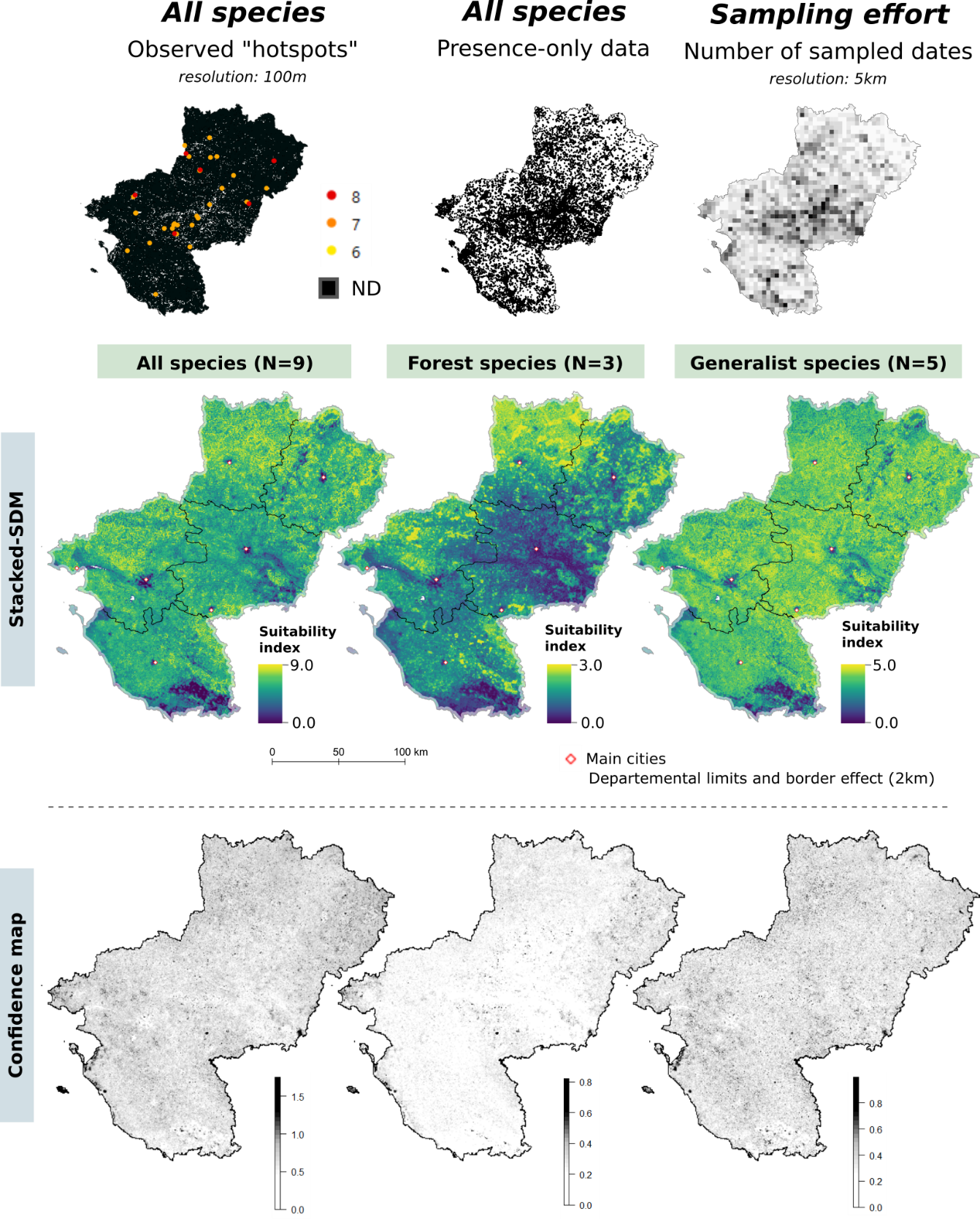

**Figure S4.1. Suitability maps from stacking species distribution models (SSDM) and associate confidence maps.** Confidence maps are the sum of individual species standard deviation map obtained by Matutini et al. 2021. Map resolution is 100m

**ALL (0.85 / 0.80 / 0.75)	FOR (0.85 / 0.80 / 0.75)	GEN (0.85 / 0.80 / 0.75)**

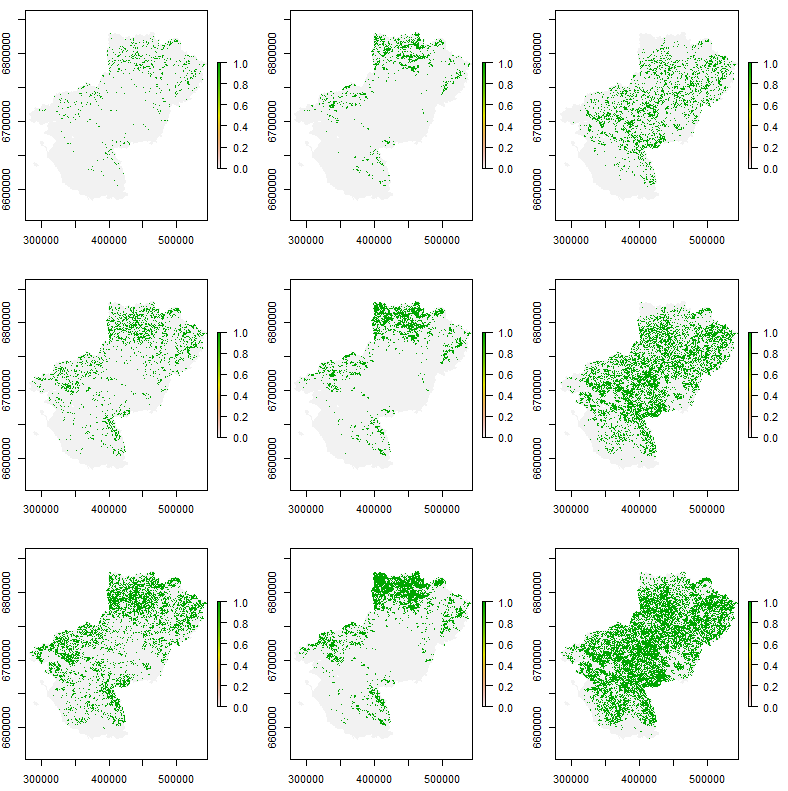

**Figure S4.2. Maps sensitivity to different thresholds selection**. Left to right (species group) : ALL, FOR and GEN ; up to down (threshold value) : 0.85, 0.80 and 0.75
