## Appendix 6 for "Conservation networks do not match the ecological requirements of amphibians"

**Appendix 6 – SCAP and future protected area projects**

The SCAP (national strategy for the creation of protected areas) was set up in 2007 and aims to strengthen the existing network by creating new protected areas to reach 2% of the PA territory. On a regional scale, these sites were defined using Atlas biological data available on the territory for 121 priority species, including 3 amphibians. In total, 174 sites were defined and 35 sites were selected in 2020 and are awaiting validation: 18 sites whose protection and / or management is to be reinforced (GR 18) as a priority by 2023, and 17 for 2030 (GR 17).

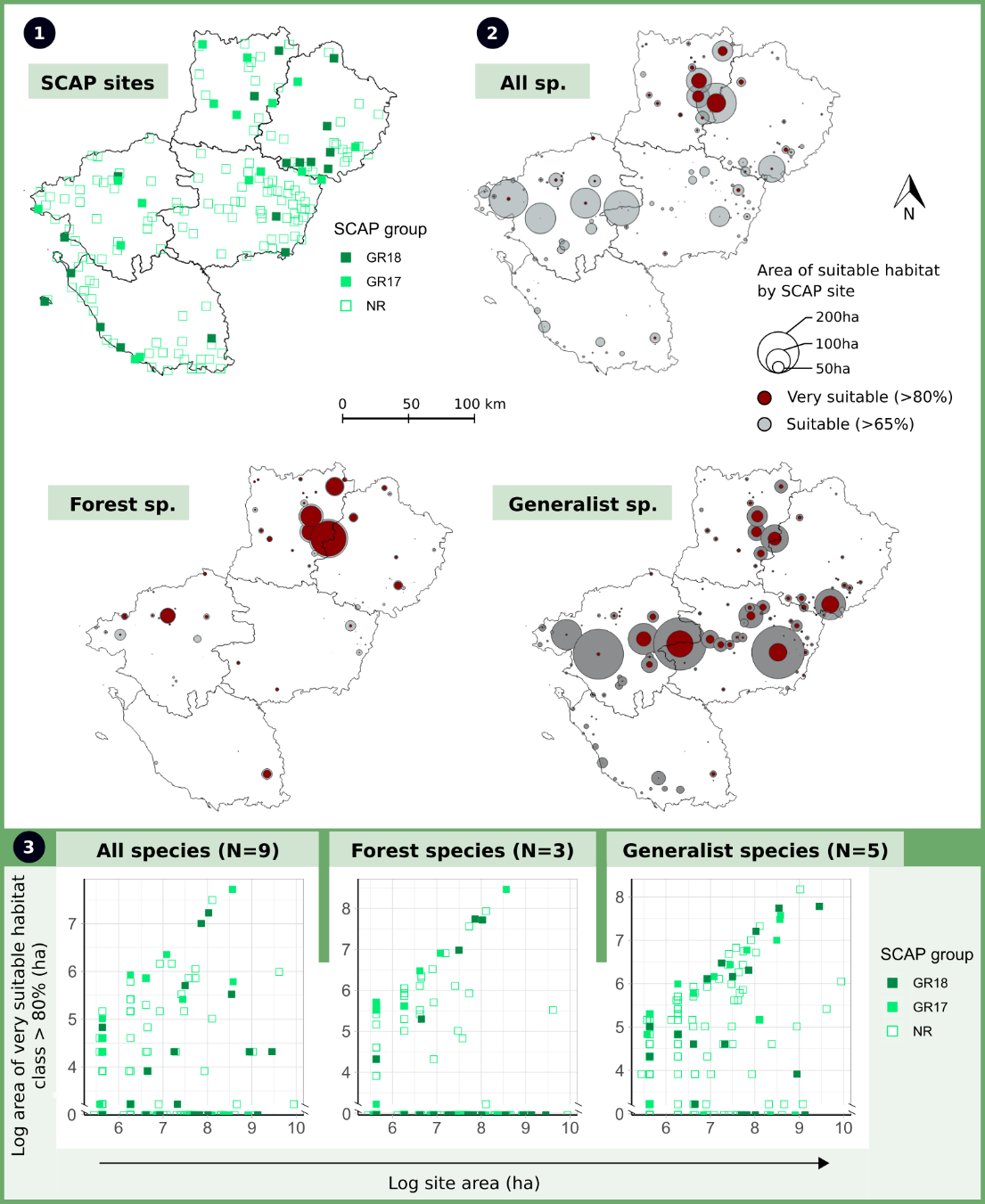

**Figure S1. Proportion of potential high suitable habitats for nine studied amphibian species sites proposed as future protected area according to SCAP strategy.**
